# Visualization and sequencing of accessible chromatin reveals cell cycle and post romidepsin treatment dynamics

**DOI:** 10.1101/2020.04.27.064691

**Authors:** Pierre-Olivier Estève, Udayakumar S. Vishnu, Hang Gyeong Chin, Sriharsa Pradhan

**Author notes:** Equal contribution (Pierre-Olivier Estève, Udayakumar S. Vishnu). Correspondence: Sriharsa Pradhan, Genome Biology Division, New England Biolabs, Inc., 240 County Road, Ipswich, MA 01938, USA.

## Abstract

Chromatin accessibility is a predictor of gene expression, cell division and cell type specificity. NicE-viewSeq (<u>Nic</u>king <u>E</u>nzyme assisted <u>view</u>ing and <u>Seq</u>uencing) allows accessible chromatin visualization and sequencing with overall lower mitochondrial DNA and duplicated sequences interference relative to ATAC-see. Using NicE-viewSeq, we interrogated the accessibility of chromatin in a cell cycle (G1, S and G2/M) - specific manner using mammalian cells. Despite DNA replication and subsequent condensation of chromatin to chromosomes, chromatin accessibility remained generally preserved with minimal subtle alterations. Genome-wide alteration of chromatin accessibility within TSS and enhancer elements gradually decreased as cells progressed from G1 to G2M, with distinct differential accessibility near consensus transcription factors sites. Inhibition of histone deacetylases promoted accessible chromatin within gene bodies, correlating with apoptotic gene expression. In addition, reduced chromatin accessibility for the MYC oncogene pathway correlated with down regulation of pertinent genes. Surprisingly, repetitive RNA loci expression remained unaltered following histone acetylation-mediated increased accessibility. Therefore, we suggest that subtle changes in chromatin accessibility is a prerequisite during cell cycle and histone deacetylase inhibitor mediated therapeutics.

## Introduction

In the human genome, genes fluctuate between various transcriptional states: active, inactive or poised (1–3). The transcriptional status of each gene is dependent on it’s local chromatin architecture and nucleosomal positioning inside the nucleus. Architectural structure of genes is dynamic during gene expression, and often dependent on external factors, including signaling molecules or internal physiological drives, such as, cell division (4–6). The local architecture of a transcriptionally active gene is characterized by accessible chromatin at the promoter, nearby enhancers, and some internal regions within the gene body itself (7–9). Biochemically these regions are also marked with active histone marks, H3K4me3 and H3K27ac that recruit the transcriptional apparatus (10, 11). For example, gene promoters are devoid of bound nucleosomes and enriched with H3K4me3 and H3K27ac nucleosomes within the vicinity that permits the binding of the transcription factors and subsequent transcriptional initiation (12–14). Therefore, it is believed that the transcriptional state of a gene is predictable by chromatin accessibility. Indeed, chromatin accessibility has become a hallmark for active DNA regulatory elements. In a study involving chromatin accessibility of 410 tumor samples, derived from 23 different primary human cancers, information on disease susceptibility, mechanism and potential therapeutic strategies was inferred from patterns of chromatin accessibility. In addition, comparison of chromatin accessibility with WGS of these patients revealed a series of somatic mutations in regulatory regions of DNA, which manifested by downstream gain of chromatin accessibility signatures and lead to poor patient prognosis (15).

Traditionally, DNase-I was predominately used to identify accessible regions (16). DNase-seq is a powerful technique for genome-wide detection of DNase hypersensitive Sites (DHS). This technique is labor intensive and requires enzyme titration to determine optimal digestion conditions for sample processing, library preparation and sequencing (17). Although a single report of single cell DNase-seq using circular carrier DNA has been reported, no other low cell input reports were published (18). Another method, FAIRE-Seq (Formaldehyde-Assisted Isolation of Regulatory Elements) has gained attention due to simplicity of accessible DNA preparation. The protocol involves sonication of cells fixed by formaldehyde to disrupt nucleosomes, followed by centrifugation (19). The accessible chromatin containing DNA remained in the supernatant and could be used for library preparation and sequencing. Other techniques focus on the ease of transposition to mark accessible chromatin regions, such as Tn5 based sequencing, ATAC-seq, as the “gold standard” of the field. Tn5 transposon-based cut and transposition (tagmentation) of the cargo DNA molecule on the accessible region has been demonstrated on native/unfixed cells. ATAC-seq (Assay for Transposase-Accessible Chromatin using sequencing) has enabled accessible chromatin studies at low cell numbers in multiple cell and tissue types (20–22). In a recent study, single cell ATAC-seq was demonstrated (23). However, due to the lower overall read coverage, data from 256 cells had to be merged to demonstrate the feasibility of the method (23). Several recent reports also demonstrate ATAC-seq’s usefulness in formaldehyde fixed cells, although the protocol demanded additional steps, including addition of DNase I to remove contaminating mitochondrial DNA, which is often overrepresented in ATAC-seq libraries. Furthermore, ATAC-seq requires Tn5 transposon titration and optimization for optimal tagmentation depending on cell type, cell numbers and treatment conditions (24). Since most clinical samples are formaldehyde fixed, Tn5 transposon based tagmentation is challenging. We have previously developed a nicking enzyme assisted sequencing (NicE-seq) method that is robust and suited to a number of cell types in native and formaldehyde crosslinking conditions (25). Currently, we have modified this method for universal adaptability by substituting dCTP with 5mdCTP and named it universal NicE-seq (Chin et al., unpublished). Here, we modified the UniNicE-seq protocol and included fluorescein-dATP or Texas Red-5-dATP in the mix to allow sequential visualization and sequencing of the accessible regions from the same sample. We named this new technology Nicking enzyme assisted viewing and sequencing (NicE-viewSeq). It is a versatile method that combines quantitative imaging and sequencing of the accessible chromatin regions across various cell type using NGS. Here we demonstrate that NicE-viewSeq can be used to investigate chromatin visualization of three-dimensional immobilized nuclei and interrogate it’s chromatin accessibility profile during cell cycle stages and post-histone deacetylase inhibitor treatment of cancer cells. We have investigated altered chromatin dynamics during cell cycle and chromatin compaction stages and post-histone deacetylase inhibitor treatment. Furthermore, we have correlated variation and alteration of chromatin dynamics with gene expression.

## Results

### Accessible chromatin visualization and sequencing

We were able to dual label accessible chromatin regions by using a deoxyribonucleotide mixture containing both biotin conjugated dCTP (bt-dCTP) and Texas-Red-dATP in the presence of DNA pol I and Nt. CviPII (Fig 1A). This is a simple one step process enabling incorporation of both fluorophore and biotin containing nucleotides. Thus, allowing us to successively visualize and perform downstream sequencing of the accessible chromatin. Indeed, the accessible chromatin in the nucleus was visualized as a red color, only in the presence of both DNA pol I, and the nicking enzyme Nt. CviPII, across three different cell lines, HCT116, K562 and GM12878. Pol I alone was not able to incorporate Texas-Red-dATP, confirming that the nicking enzyme is an essential component for visualization (Fig 1B). We were also able to superimpose our accessible chromatin signal on a nuclear DAPI staining pattern demonstrating that accessible chromatin and heterochromatin are mutually exclusive, and also, showing that the mitochondrial DNA signal was not detectable (Supp Fig. 1). We extracted DNA from HCT116 cells and captured the biotin and Texas-Red incorporated accessible chromatin on streptavidin magnetic beads and constructed a library for NicE-viewSeq (Fig 1A). The NicE-viewSeq library displayed a similar accessible chromatin pattern when compared with stand-alone accessible chromatin sequencing profiles derived from NicE-seq, DNase-Seq and ATAC-seq of the HCT116 cell genome (Fig 1C). We further compared NicE-viewSeq and UniNicE-seq of HCT116 using Pearson correlation of peak read densities and observed a correlation value of r=0.98 (Fig 1D). A similarly high degree of correlation was also observed when ATAC-seq and NicE-viewSeq were compared (r=0.83), demonstrating that the global picture of accessible regions is common between NicE-viewSeq, NicE-seq and ATAC-seq (Supp Fig 2A). The TSS profile between NicE-viewSeq and universal NicE-seq also remain similar, demonstrating addition of fluorophore containing nucleotides to the reaction does not alter capture and analysis of TSS (Fig. 1E). Furthermore, we also used HeLa cells to perform NicE-viewSeq and compared it with the accessible chromatin profiles of ATAC-seq and DNase-seq. As expected, we observed similar accessible chromatin profiles in all three different techniques (Supp Fig 2B). An overall strong accessible chromatin correlation was observed between ATAC-seq and NicE-viewSeq (r=0.6) along with similar TSS metaplots (Supp Fig 2C-D). Taken together these results, we demonstrate that accessible chromatin can be labeled by both fluorophore and biotin conjugated dNTPS simultaneously for viewing, quantitative imaging and NGS applications without mitochondrial DNA background.

**Figure 1:**
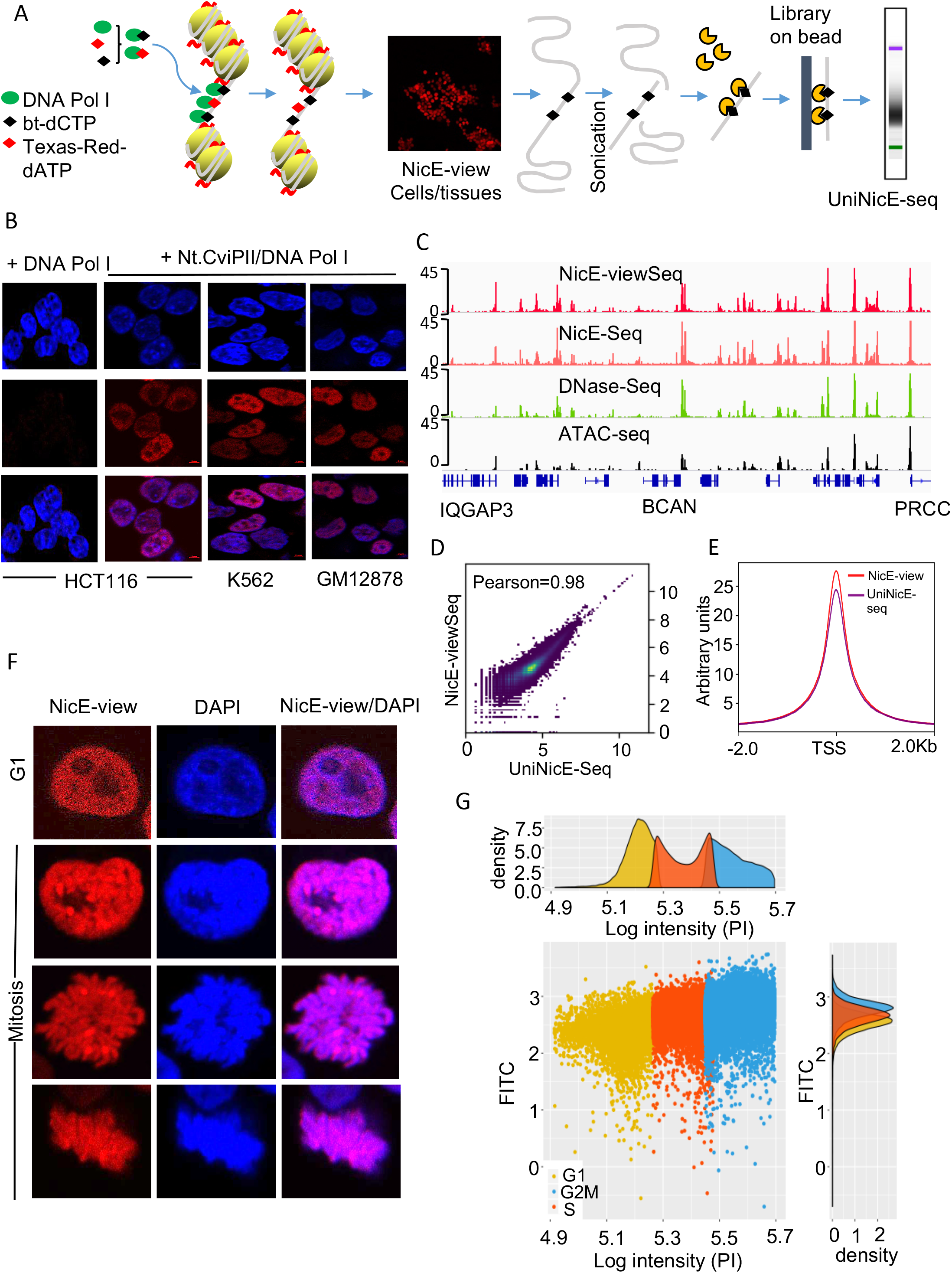
NicE-viewSeq optimization and validation. (A) Schematic diagram of NicE-viewSeq method for visualization and NGS library preparation. (B) NicE-viewSeq labeling in fixed HCT116, K562 and GM12878 cells using dNTPs supplemented with Texas Red–dATP. Top panel (DAPI), middle panel (Texas Red) and lower panel (merged). (C) IGV genomic tracks of accessible chromatin using NicE-viewSeq and comparison with NicE-seq, DNase-Seq and ATAC-seq. Gene names shown at the bottom. (D) Genome-wide comparison of accessible chromatin between universal NicE-seq (UniNicE-seq) and NicE-view Seq using sequence read density. (E) Metagene plot based on transcription start site (TSS) between universal NicE-seq (UniNicE-seq) and NicE-view Seq. (F) Representative NicE-viewSeq of cycling cell during cell cycle depicting G1 and mitotic phase. Please note that mitotic cells have 4n DNA and more brightly stained compared to G1 (2n) cell. (G) Scatter and density plot showing G1, S and G2M phases of cell cycle based on PI fluorescence. The fluorescence increases from G1 to G2M phase.

Hyperactive Tn5 transposon mediated ATAC-see offers a convenient cut and tag method of inserting fluorophore labelled DNA into mammalian cells for visualization and sequencing of accessible chromatin. We therefore compared accessible chromatin data and method specific visual changes between ATAC-see with NicE-viewSeq of HT1080 cells. Although the FRiP scores between both methods were comparable between 0.25-0.30, mitochondrial reads in ATAC-seq were 65%, compared to the lower 14% in NicE-viewSeq. Similarly, percent duplication of reads in ATAC-see were relatively higher in HT1080 (66-70%) and GM12878 (93-97%), compared to NicE-viewSeq in HT1080 (13-15%), HUT78 (7-16%) and HCT116 (39-49%) (Supp table 2A, 2B, 3, 4, 8). Visualization of accessible chromatin peaks were similar in both methodologies, although Pearson’s correlation of either all the mapped reads or read density of the peaks between NicE-viewSeq and ATAC-see for HT1080 fixed cells were r=0.6 compared to unfixed HT1080, r=0.45 (Supp Fig 3C). These results suggest that NicE-viewSeq enables higher accessible chromatin peak calling at lower sequencing depth with relatively lower mitochondrial read and duplicated sequences count.

### NicE-viewSeq of accessible chromatin during cell cycle and sorting

Since NicE-viewSeq is a simpler method without the need for enzyme to cell number titration, we used it in cell cycle studies. Nuclear chromatin undergoes distinctive structural alterations during various cell cycle stages, where both compaction and decompaction of chromatin takes place. Indeed, in the G1 stage, the cell is metabolically active and chromatin is in a stable state; S phase partially condenses chromatin and thereby, facilitates DNA replication; G2 and M (mitosis) phase brings chromatin compaction. Therefore, we hypothesized that cell-cycle specific changes of chromatin accessibility can be studied using NicE-viewSeq to identify dynamic accessible chromatin domains during cell cycle; as shown for HeLa cells using accessible chromatin labeling, supplemented with Texas-Red conjugated dATP. We noticed accessible chromatin labeling at G1 stage is distributed throughout the nucleus with a small punctate pattern. The mitotic cells also displayed accessible chromatin labeling with a more distinctive localized signal suggesting preservation of accessible chromatin during mitosis (Fig 1F). Some of this increase in the accessible chromatin signal would be attributed to duplication of DNA between G1 to M stage from 2n to 4n. To perform NicE-viewSeq labeling and FACS sorting of HCT116, we used fluorescein-dATP along with biotinylated-dCTP and further stained the cells with PI (propidium iodide). We collected cells from different cell cycle stages using an established protocol (26). Fluorescein-dATP labeling was used as an internal control for visual confirmation of accessible chromatin labeling, and the percentage of cells in G1, G2M and S phase of the cell cycle were determined by total cellular DNA content analysis using propidium iodide exhibiting higher accessible chromatin labeling as the total gDNA content increases between G1 to G2M (Fig 1G, Supp Table 1). Between 200,000 and 500,000 cells were collected for subsequent accessible chromatin analysis.

### Dynamic accessible chromatin landscape during cell cycle

Since the cells were labeled with fluorescein-dATP and biotin-dCTP before FACS sorting, each fraction representing G1, G2/M and S cell populations, were used directly for DNA extraction and NicE-seq library preparation. All NicE-seq reads were downsized to 13 million and we obtained 53, 43 and 26K accessible chromatin peaks for G1, S and G2M cells, respectively (Supp Table 3). The NicE-viewSeq based chromatin accessibility profile of HCT116 cells often coincides with promoters and vicinity of transcription factors. Chromatin accessibility often is a predictor of gene expression. To demonstrate this phenonemon, we singled out three well-studied cytosine-5 DNA methyltransferase genes, DNMT1, DNMT3A and DNMT3B, which participate in epigenetic inheritance and examined their chromatin accessibility during cell division in human cells. The IGV browser displayed preserved chromatin accessibility for G1, S and G2/M cells, coinciding with H3K27ac ChIP-seq and DNase-seq peaks, displaying both accessible promoters and intergenic regions for all three DNA methyltransferases (Fig 2A). However, DNMT1, the major maintenance methyltransferase that methylates newly synthesized daughter strands during DNA replication during the S phase of cell cycle, displayed a significant reduction in promoter chromatin accessibility as the cell exists S phase and enters G2/M phase. Both *de novo* methyltransferases, DNMT3A and DNMT3B, chromatin accessibility during G2M phase were also reduced but not as prominently as DNMT1. These observations demonstrate dynamic chromatin accessibility during cell cycle that impacts epigenetic inheritance via DNA methylation. We also examined accessible chromatin of genes periodically expressed in human cell cycle, G1/G1S, S and G2/M (27). In general, at the TSS (−2 to +2kb), accessible chromatin of displayed a higher sequence read density at G1/G1S and S, compared to the G2/M stage (Fig 2B). When tag densities of all accessible peaks for TSS and enhancers were compared, G1 cells represented the highest, followed by S and G2M cells (Fig 3A-B). This would imply that the chromatin accessibility of TSS and enhancer elements are gradually decreased as the cell cycle progresses. Taken together this would confirm that accessible TSS and enhancers are present throughout the cell cycle and their dynamic accessibility may regulate transcription during the DNA compaction and mitosis.

**Figure 2.**
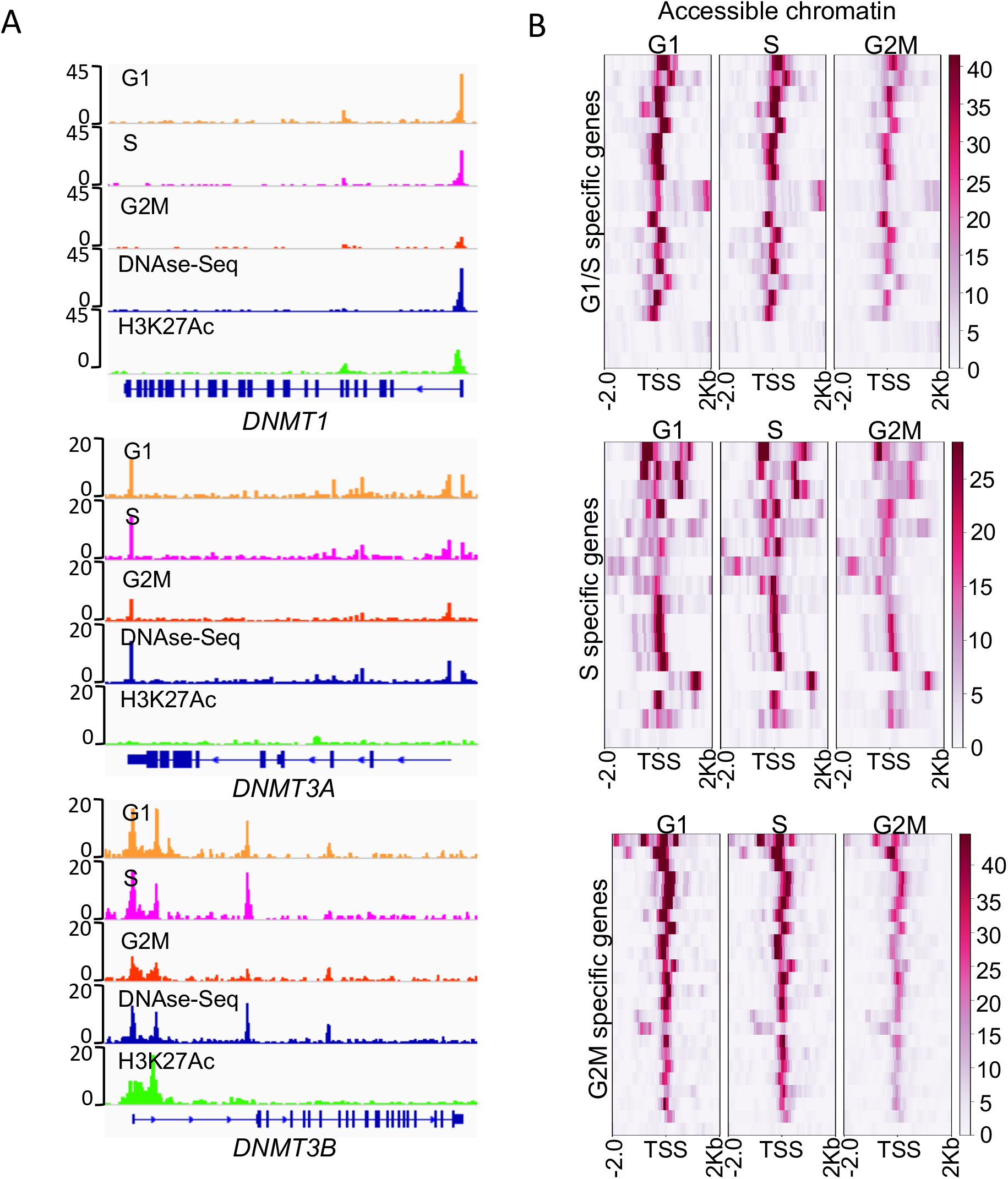
Accessible chromatin and cellular localization of DNA methyltransferases during cell cycle. (A) IGV genomic tracks of accessible chromatin using NicE-viewSeq and correlation with DNase -seq and H3K27 genomic features. DNA cytosine-5 methyltransferase 1 (DNMT1), DNA cytosine-5 methyltransferase 3A (DNMT3A) and DNA cytosine-5 methyltransferase 3B (DNMT3B) are shown. (B) Heat map of accessible chromatin of TSS regions of known cell cycle regulated genes at G1/G1S, S, and G2M stages.

To determine the relative similarities of accessible chromatin between different cell cycle phases, we performed Pearson’s correlation analysis using peak and tag densities of reads as the main criteria. Indeed, G1 and S cycle cells displayed a high degree of correlation with r=0.89 compared to G1 vs. G2/M (r=0.41) and S vs. G2/M (r=0.69), suggesting a high degree of transitional overlap between accessible regions as the cell enters G1 to S (Fig 3C (read density at peak correlation), Supp Fig 4A (mapped reads correlation)). We also observed a set of sequence reads in G2/M that did not correlate either with G1 or S phase. To negate this as an experimental artifact, we also treated cells with nocodazole to arrest them in G2/M and then compared nocodazole reads with G1 and S phase reads; we obtained similar sequence read density correlations (G1 vs. S r=0.91; S vs. G2M nocodazole r=0.66; G1 vs. G2M nocodazole r=0.52; Supp Fig 4B). Furthermore, we also compared cell cycle accessible chromatin analysis of previously published suspension cells GM12878 ATAC-seq datasets and observed a similar pattern mapped read correlation of accessible chromatin (GM12878: G2M r=0.77; HCT116: r=0.44 and 0.52). Mapped reads correlation between G1 and S were more prominent in HCT116 cells compared to GM12878 ATAC-seq datasets (HCT116: G1 vs. S r=0.94 and 0.91; GM12878: r=0.60) and the converse was true for correlation between S to G2/M suggesting cell type and cell division stage specificity (HCT116: S vs. G2M r=0.64 and 0.66; GM12878: r=0.94; Supp Fig 4A-C) (28). Based on all three different experimental analysis, we confirm that G2/M phase of the cell-cycle brings on unique sets of accessible chromatins compared to G1 and S phase.

**Figure 3.**
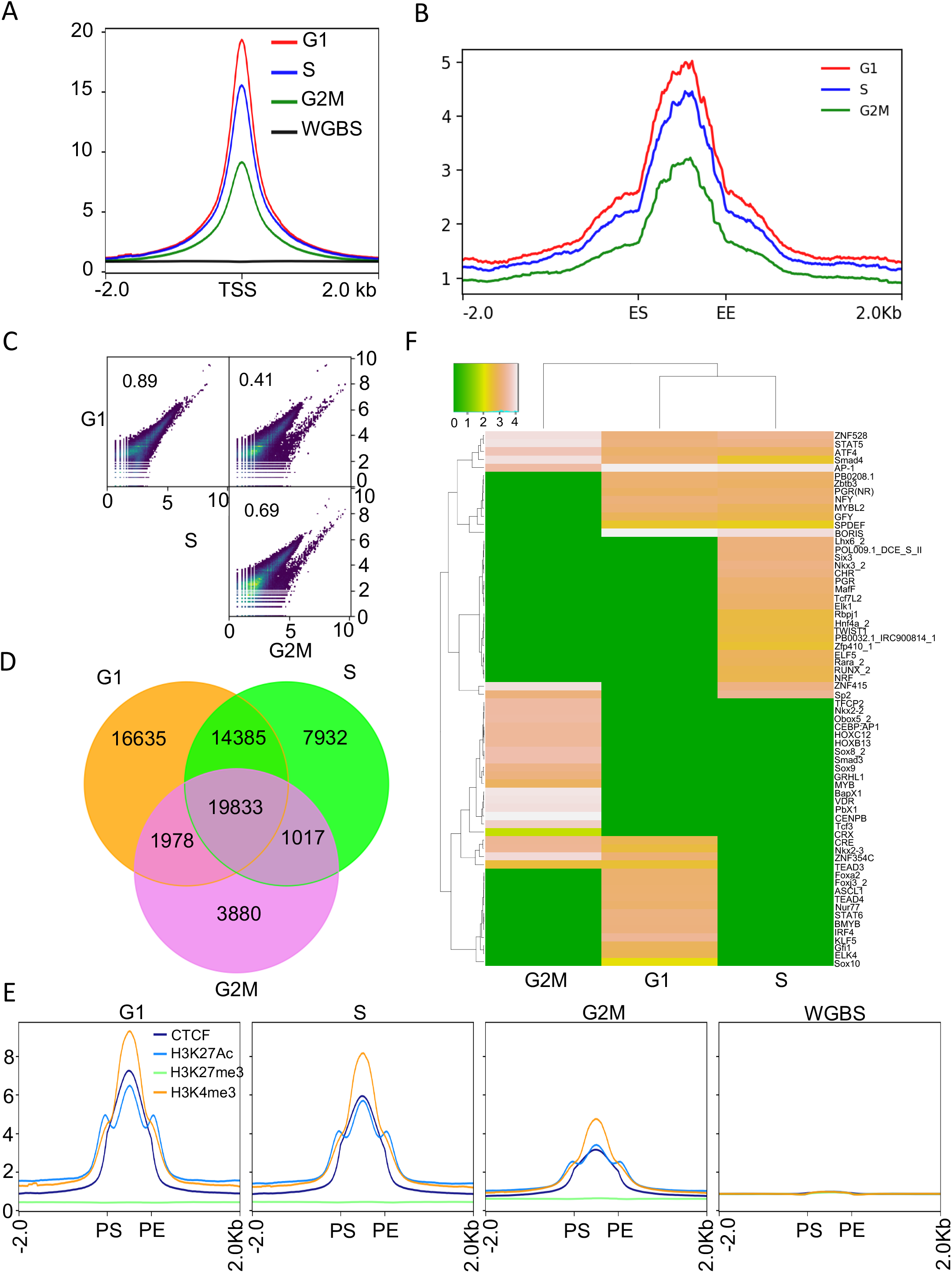
Genome wide accessible chromatin analysis during HCT116 cell cycle using NicE-viewSeq. (A) Genome-wide metagene plot of TSS with ± 2Kb of flanking region of NicE-viewSeq analysis along with corresponding control whole genome bisulphite (WGBS) analysis. (B) Genome-wide metagene plot of enhancer elements with ± 2Kb of flanking region following enhancer start (ES) and enhancer end (EE) site of NicE-viewSeq analysis. (C) Pearson correlation between accessible region reads of G1, S and G2M phases using scatter plot display. R values are inserted and shown. (D) Venn diagram showing common and unique accessible region peaks between G1, S and G2M phases. (E) Signal intensity profile plot showing the enrichment of CTCF, H3K27Ac, H3K27me3 and H3K4me3 in NicE-viewSeq analysis of G1, S, and G2M phases along with WGBS as control. Peak-start and peak-end are noted as PS and PE. (F) Heatmap showing transcription factor binding consensus motifs enrichment near open-chromatin regions corresponding to NicE-viewSeq.

Next, we called all accessible peaks in each cell cycle stage and compared them. Although, a majority of accessible peaks were common during cell cycle stage, a significant percentage of accessible regions were unique to a specific cell cycle stage (Fig 3D). Furthermore, we also extracted all accessible promoter peaks from G1, S and G2M stages and analyzed the correlation between all cell-cycle specific peaks. The Pearson’s correlation between G1 vs. S (r=1), S vs. G2M (r=0.94) and G1 vs. G2M (0.96) demonstrated strong correlation (Supp Fig 4D). This suggested that cell-cycle stage specific accessibility of the chromatin is preserved in most of the promoter regions. We further analyzed these regions in order to compare if accessibility is correlated with canonical DNA or histone modification signatures, since these epigenetic marks are dynamic and establish cell fate. For this analysis, we compared G1, S, G2/M specific accessible chromatin regions with two active histone marks, namely H3K27Ac and H3K4me3, alone with, one inactive histone mark H3K27me3, and lastly, CTCF transcription factor binding regions of asynchronous HCT116 cells that were obtained by ChIP-seq. Indeed, we observed a strong correlation of accessible chromatin with active histone marks and CTCF binding sites, and a summarily poor correlation with inactive histone marks and DNA methylation as expected (Fig 3E). These results show dynamic accessible chromatin regions during cell cycle are marked by canonically active histone modifications.

### Cell cycle specific transcription factor sequence enrichment in accessible chromatin vicinity

During cell division a variety of sequence-specific transcription factors regulate gene expression by binding to cis-regulatory elements in promoter and enhancer DNA. Since promoter and enhancer elements are accessible during transcription, we hypothesized that NicE-seq could be a predictor of transcription factor binding. To validate our hypothesis, we performed NicE-viewSeq sequence read density comparison around accessible chromatin peaks (−2 to +0.1 kb) between G1, S and G/2M cells and investigated the proximity of consensus transcription factor binding sequences to accessible chromatin peaks during cell cycle. We observed varying degrees of similarity and contrast between accessible chromatin between G1, S and G2/M cells demonstrating cell-cycle specific transcriptional programming (Fig 3F). For example, sequence read density enrichment of RUNX2 binding site was prominent in S phase, coinciding with the previous observation of increased RUNX2 protein levels during G1/S (29), and G1 to S entry (30). We also observed enrichment of KLF5 (Kruppel-like factor 5), STAT6 and BMYB consensus sequences close to accessible chromatin during G1 phase. KLF5 is a basic transcription factor that binds to GC boxes at a number of gene promoters and is involved in transcriptional of those genes during development. In adults, KLF5 expression is higher in proliferating epithelial cells (31). Similarly, the presence of STAT6 signature enrichment in G1 also suggests that it may assist cell cycle dependent enhanceosome at target gene promoters by recruiting transcriptional enhancers. Indeed, CBP/p300 is recruited by STAT6, and this association results in increased STAT6-dependent transcription (32). BMYB transcription factor is known to promote Saos-2 cells into the S phase of the cell cycle and to overcome G1 arrest (33). Other cell cycle specific transcription factors sequences, particularly for G2/M such as CENBP, VDR and TFCP2 (LSF) were also enriched. While CENBP is a major centromere formation protein during mitosis, both VDR and LSF play active role in G2/M phase. Therefore, these results demonstrate the versatility of NicE-viewSeq, where cells could be labeled or stained for primary visual analysis followed by downstream genomic analysis.

### NicE-view (Nicking Enzyme assisted viewing) for global chromatin accessibility post epigenetic drug treatment

Since accessible chromatin could be fluorescently labelled using fluorophore-dATP, we hypothesized that alteration of genome architecture by epigenetic drugs could be monitored in a dose dependent manner. For proof of concept, we used HUT 78 cells, a T cell lymphoma cell line, and treated those cells were with romidepsin, an FDA approved therapeutics for T cell lymphoma. Romidepsin is a prodrug with the disulfide bond that undergoes reduction in the cell and releases a zinc-binding thiol moiety. The thiol moiety binds to a zinc of histone deacetylase and blocks it’s enzymatic activity; thus acting as a histone deacetylase inhibitor (HDACi). Inhibition of HDAC would promote histone acetylation, and thus would increase chromatin accessibility. We monitored HUT 78 cells after romidepsin treatment using NicE-view (Fig 4A), and also performed western blot analyses for the H3K27acetyl mark to demonstrate its efficacy (Fig 4B). Indeed, H3K27 acetylation increased 6-fold following 6 hrs romidepsin treatment compared to control (Supp Fig 5A). We also measured the NicE-view pixel intensity and observed genome-wide a 2-fold gain of chromatin accessibility with a p-value <0.001 (Fig 4B). This observation demonstrates that NicE-view can be used for chromatin accessibility quantitation post epigenetic therapy using cell-based assay.

**Figure 4.**
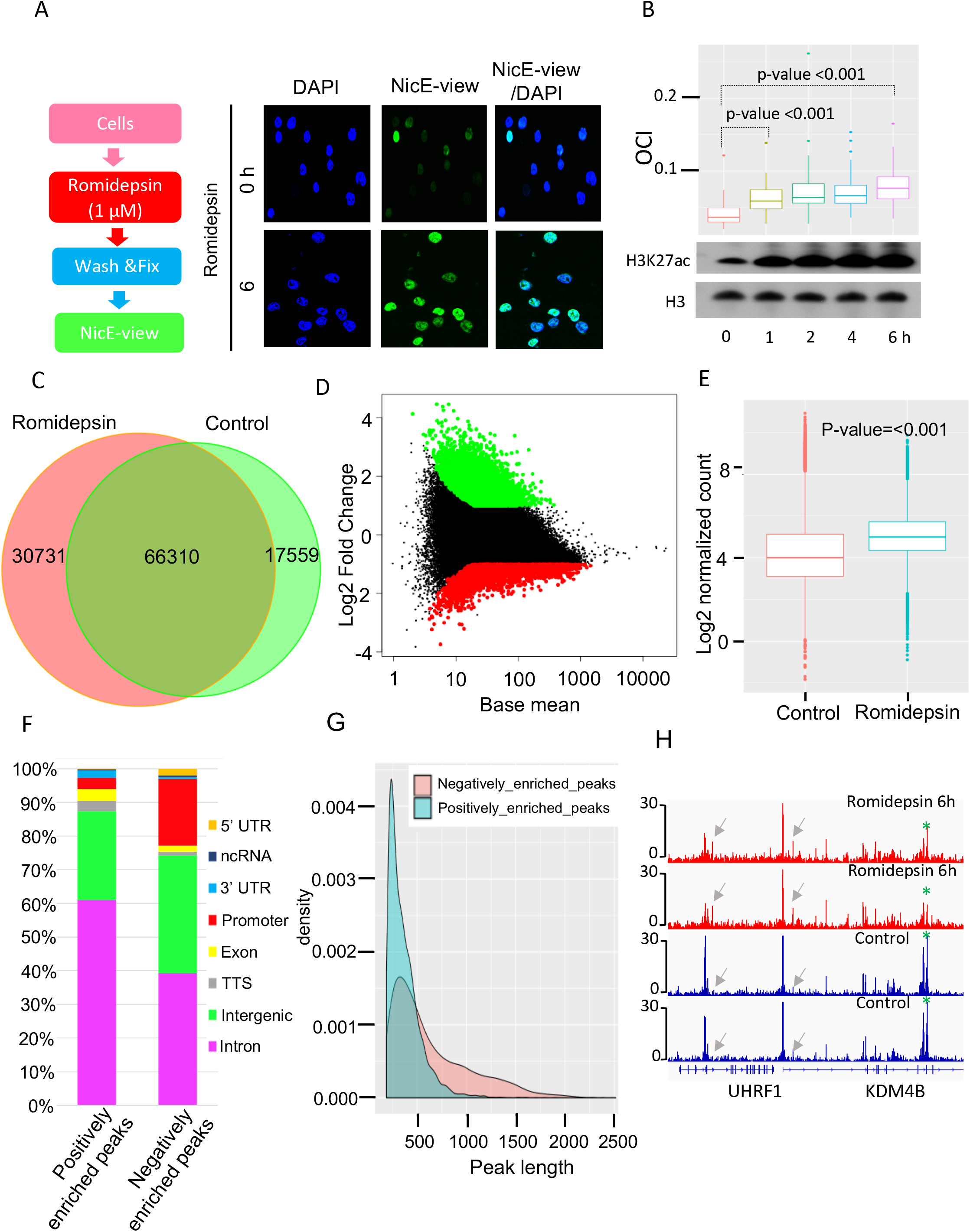
HDAC inhibitor romidepsin and accessible chromatin analysis using NicE-viewSeq using HUT 78 cells. (A) Accessible chromatin labeling and visualization scheme on left and images on right at 0 and 6 hr post drug treatment. Merged images shows nuclear staining and NicE-viewSeq staining. (B) Time course of romidepsin treatment and nuclear accessible chromatin quantitation as indicated by open chromatin index (OCI) on the top and western blot for H3K27ac and H3 at the bottom. OCI was calculated as a ratio between pixel density of the fluorophore and DAPI. (C) Venn diagram showing common and unique peaks between control and romidepsin treated HUT 78 cells. (D) MA plot showing the positive and negatively enriched accessible chromatin peaks after the romidepsin treatment. (E) Boxplot (the line the box shows the median) of control and romidepsin treated HUT 78 cells showing more read counts in treated cell indicating more accessible chromatin after the romidepsin treatment. (F) Peak annotation of positive and negatively enriched peaks after the romidepsin treatment showing the distribution of peaks. (G) Density plot based on the peak length of positive and negatively enriched peaks after the romidepsin treatment. (H) IGV Genomic tracks of differential accessible chromatin at 0 and 6 h post romidepsin in experimental duplicates. Reduced or gain of accessibility is shown at specific locations, although genome wide accessibility was increased after romidepsin treatment.

### DNA accessibility landscape post romidepsin treatment

To identify the difference in dynamic chromatin accessibility, we performed universal NicE-seq libraries of both control and romidepsin treated samples and sequenced in depth (Supp Table 4). Chromatin accessibility throughout the genome increased after 6 hrs of romidepsin treatment, including appearance of newly acquired accessible regions (Fig 4C). Hierarchical clustering of two biological replicates, each with two technical replicates for control and romidepsin treated cells, displayed a high degree of correlation of accessible chromatin regions (Supp Fig 5B). The principal component analysis of both data sets showed a high degree of correlation among experimental duplicates but distinct changes in romidepsin treated cells compared to control (Supp Fig 5C). Indeed, accessible chromatin density increased, and length decreased in romidepsin treated cells (Supp Fig 5D). Most of the promoters after romidepsin treatment displayed reduced promoter accessibility (Supp Fig 5D). Overlapping accessible chromatin peaks were 70-80% between both samples suggesting the majority of the accessible regions did not change (Fig 4C). However, 21% of accessible regions were exclusive to romidepsin treated cells, demonstrating HDACi specific accessible chromatin peaks. This was further supported by differential analysis of NicE-seq read count data per accessible peak (Fig 4D), using shrinkage estimation for dispersions and fold changes (Love et al. 2014). We finally performed log2 normalized counts for control and romidepsin treated peaks and observed that, as expected, HDACi increased the quantity of accessible chromatin peaks (Fig 4E). Furthermore, most positively enriched accessible chromatins were situated in the introns and negatively enriched peaks at promoters suggesting alteration of chromatin accessibility occurs predominately within introns and promoters (Fig 4F). However, positively enriched peaks were narrower compared to broad negatively enriched peaks that constitute the promoters, suggesting romidepsin can adversely affect gene expression via promoter accessibility alteration (Fig 4G-H).

### Change in chromatin accessibility and differential gene expression

We further examined if the differential accessible chromatin is correlated with transcriptome profiles. We performed RNA-seq of both control and 6-hr romidepsin treated cells (Supp Table 5). Hierarchical clustering of three technical replicates of control and romidepsin treated cells displayed a high degree of correlation of expressed RNA (Supp Fig 6A). The volcano plot demonstrated that almost an equal number of differentially expressed genes were both positively and negatively regulated, similar to strong signature of loss or gain of accessible chromatin regions post-romidepsin treatment (Fig 5A, Fig 4C, Fig 4D, Supp table 6). The genes associated with the differentially enriched peaks from NicE-seq were identified and compared with the differentially expressed genes identified from RNA-seq. About 60% of the accessible DNA were unique and ~17% correlate with RNA expression (Fig 5B). About 25% transcriptionally active RNA did not overlap with any accessible chromatin region. When tag densities between control and romidepsin RNA seq and NicE-seq were compared, there was a small but significantly higher sequence read density in romidepsin treated cells for accessible chromatin and gene expression. For example, NicE-seq control and NicE-seq romidepsin vs. RNA-seq romidepsin tag has an increased correlation from 0.59 to 0.67. Therefore, the fold changes in accessible chromatin and transcription is consistent (Fig. 5C). We narrowed down our accessible chromatin regions with specific gene expression. For this study, we interrogated the top 200 differentially accessible chromatin regions with gene expression and observed 4 different patterns. Correlation of accessible chromatin regions was observed depending on the group. In group I high accessibility of chromatin correlated with gene expression, whereas low accessibility regions correlated with lower gene expression as shown in group IV. However, in group II lower accessible chromatin displayed higher gene expression and the inverse was true for group III (Fig. 5D, Supp table 7). These data suggest that although there is a direct correlation between chromatin accessibility and gene expression, a subset of genes do not follow that path and may be regulated by a different mechanism. Indeed, repetitive DNA elements such as LINE, SINE and LTR gained chromatin accessibility without transcriptional activation (Supp Fig 6B, C). This observation is in line with a previously published report showing hypermethylated genes cannot be transcriptionally reactivated with HDAC inhibitor alone in tumor cells, but require DNA methylation inhibitor in combination with HDACi (34).

**Figure 5.**
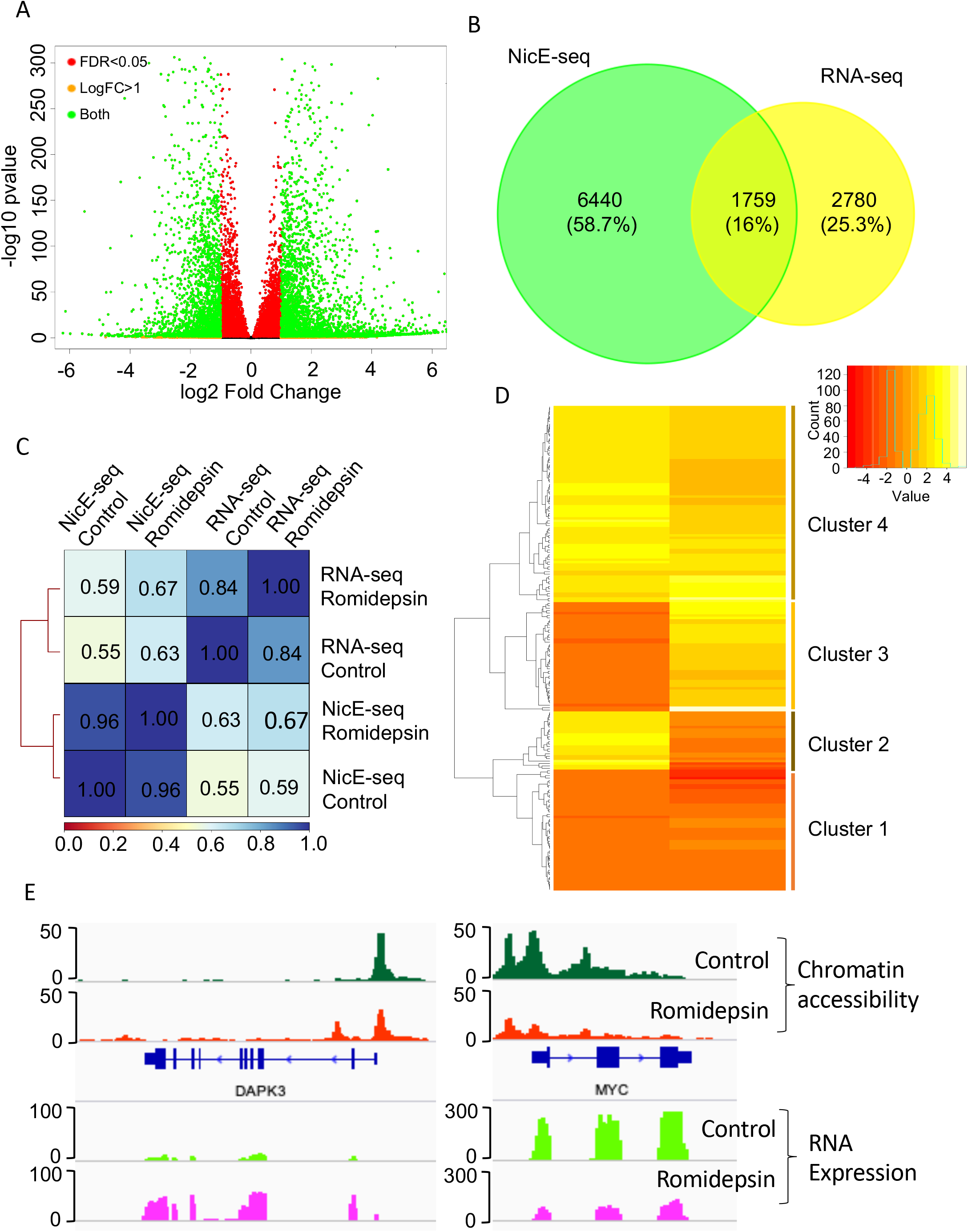
Romidepsin treated transcriptional alteration and chromatin accessibility using RNA-seq and NicE-seq. (A) Volcano plot showing the differentially expressed genes after the romidepsin treatment (logFC≥1 and FDR<0.05). (B) Venn diagram showing the congruence between genes associated with differential accessible chromatin regions (peaks) and differentially expressed genes from RNA-seq. (C) Spearman correlation between NicE-Seq and RNA-seq using tag densities. The correlation (r) values are shown. (D) Heatmap of top two hundred genes showing the congruence between the differential accessibility in the gene body and differential expression in RNA seq based on their fold change. (E) Normalized NicE-seq profiles of control and romidepsin treated cells in DAPK3 and MYC locus showing more accessibility in apoptotic related genes and reduced accessibility in MYC oncogene. Their corresponding gene expression post romidepsin treatment were mentioned on the top.

### Regulatory pathways gene expression partially correlated with accessible chromatin

In response to romidepsin, the apoptotic pathways are activated (Supp Fig 7), and the phosphatidylinositol 3-kinase/AKT/mammalian target of rapamycin and β-catenin pro-survival pathways were similarly inhibited (Supp Table 9). We investigated if the accessible chromatin regions of genes involved in the above pathways correlated with their expression level. Indeed, the pathway genes fall in two different categories. In the 1^st^ category, with response to romidepsin the promoter accessibility decreased, and gene body accessibility increased, resulting in higher gene expression, as observed for DAPK3 (1.83 folds higher expression). In the second category, the promoter accessibility and gene body accessibility decreased, resulting in reduced gene expression, as observed for MYC (−2.01 folds lower expression) (Fig 5E). Similarly, the highest level of RNA expression was observed for key apoptotic genes such as SERPINB9 and MAPK8IP1 (Supp Fig 7).

## Discussion

Here we demonstrate a novel genome-wide accessible chromatin visualization, quantitation and sequencing method that enables deciphering of alterations within genome hierarchy during cell cycle stages and following epigenetic drug treatment in human cultured cells. We uncover several genomic features that are specific to G1, S and G2M phases of the cell-cycle. Firstly, reduction of chromatin accessibility was the main feature when cells entered from S to G2/M phase. We also observed that accessible chromatin regions during cell-cycle are preserved and dynamic throughout. In another study, DNase-seq analysis of murine erythroid cells at different mitotic stages revealed a preserved chromatin accessible pattern (35). In our study, decrease of relative accessibility of chromatin suggests that cell cycle specific protein factors may have a role in chromatin organization. This was supported by differential transcription factor consensus binding sequence during G1, S and G2/M. Indeed, G2/M-specific transcription factors may evict nucleosomes from DNA and chromatin, remodel the surrounding architecture to recruit other proteins for transcriptional activity, thereby increasing the accessibility of specific regions. The transcription factor enrichment sequences during cell cycle phase, and gene ontology analysis, displayed cell cycle dependent signaling pathways. In addition, restart of transcription after DNA replication is required to establish chromatin accessibility (36). Thus, NicE-viewSeq can be adapted to quantitative imaging of the cells, as well as, NGS based sequencing methodologies of their genomes. Another Tn5 transposon-based Assay for Transposase Accessible Chromatin with Visualization (ATAC-see) that adds fluorescent clusters together with DNA markers has been reported. ATAC-see allows visualization of immobilized nuclei along with strong mitochondrial signal due to preferential tagmentation of mitochondrial DNA (28). Indeed, ATAC-see image processing workflow requires masking of strong mitochondrial signal to visualize nuclear signal of the cell. Compared with this method, NicE-viewSeq background signals were minimal and the nuclear signal more prominent. Sequence analysis of ATAC-see samples for accessible chromatin capture and sequencing represented ~5 fold excess of mitochondrial DNA sequence reads compared to NicE-viewSeq of fixed HT1080 cells demonstrating lower mitochondrial burden (Supp Table 2A vs. 2B). NicE-viewSeq performed on fixed HeLa and HUT 78 cells represented 6 – 15% of the mitochondrial reads (Supp 4). These results demonstrate the ease and versatility of NicE-viewSeq.

A fundamental drawback in clinical investigations and diagnostics is the use of different sections of biological material for visualization and molecular analysis, therefore disconnecting the spatial organization with molecular analysis. Our data indicates that both visualization and sequencing for open chromatin could be performed in parallel within the same sample. Here we show the proof of principle using several different cell lines. NicE-viewSeq offers spatial organization of the accessible chromatin in the nucleus for any cell type by incorporating fluorophore and biotin conjugated-dNTP and is amenable to flow cytometry. The simplicity of NicE-viewSeq on fixed cells makes it readily adapted to human clinical samples where morphology/immune histochemistry based cellular identification can be followed by chromatin accessibility studies to determine alteration of disease pathways by following marker and accessible genes. Another potential application of NicE-viewSeq is in epigenetic drug screening studies. Due to the recent developments in chromatin biology, novel drugs have been identified which are directed at chromatin and associated components, particularly DNA and histone modifications. Any impact on DNA methylation, histone modification, particularly acetylation/deacetylation, and methylation will have a direct impact on chromatin accessibility. Cell based assays involving quantitation of chromatin accessibility and gene expression/alteration will contribute to determining efficacy and cellular physiology. In this study HUT 78 cells treated with romidepsin not only showed genome-wide increase in quantitative chromatin accessibility, but also activation of specific apoptotic pathways and related genes. In summary, NicE-viewSeq greatly facilitates quantitative imaging and epigenomic analyses in both research and possibly in clinical settings.

## Materials and methods

### Open chromatin labeling with fluorescent dNTPs

HCT116, HeLa or HeLa-S3 and HT1080 cells were grown on slides. Cells were cross-linked using 1% formaldehyde for 10 min at room temperature. Formaldehyde was quenched by 125 mM glycine. Cytoplasm was extracted by incubating the cross-linked cells in cytosolic buffer (15 mM Tris-HCl pH 7.5, 5 mM MgCl_2_, 60 mM KCl, 0.5 mM DTT, 15 mM NaCl, 300 mM sucrose and 1% NP40) for 10 min on ice with occasional agitation. Slides were then washed twice with the cytosolic buffer. Fluorescent open chromatin DNA labeling was performed by incubating the nuclei in presence of 2.5 U of Nt.CviPII (NEB R0626S), 50 U of DNA polymerase I (M0209S) and 30 μM of each dNTP including 6 μM of Fluorescein-12-dATP (Perkin Elmer, NEL465001EA) or 6 μM of Texas Red-5-dATP (Perkin Elmer, NEL471001EA) in 800 μl of 1 × NEBuffer 2 and carried out at 37°C for 2 h. 80 μl of 0.5 M EDTA and 2 μg of RNase A was added to the labeling reaction and incubated at 37°C for 30 min to stop the labeling reaction and digest cellular RNA. After washing the slides once with PBS, a subsequent wash with PBS including 0.01% SDS and 50 mM EDTA was performed at 55°C for 15 min to remove fluorescent background followed by 3 washes with PBS for 5 min at RT. Slides were dried, mounted using Prolong Gold antifade reagent with DAPI (Invitrogen, P36935) and visualized using LSM 880 confocal microscope (Zeiss). Fluorescein-dATP, TexasRed-dATP and DAPI were detected using Argon 458,488,514 nm, DPSS 561 nm and diode 405 nm laser respectively. Quantification of Fluorescein-dATP or Texas Red-dATP per nucleus was defined by mean pixel intensity per nucleus using the histogram and colocalization tools included in the Zen software (Zeiss). For HUT 78 suspension cells, DNA crosslinking with 1% formaldehyde was performed in Eppendorf tube in 1 ml total volume reaction. Nuclei were labeled in 800 μl of 1 x NEBuffer 2 with constant rotation at 37°C for 2 h. After blocking the reaction with EDTA, cells were spun down at 1000 rpm for 10 min on slides and crosslinked with 1% formaldehyde for 10 min. Slides were then washed at 55°C as described above. We followed the above protocol for K562 and GM12878 cells fixation.

### Open chromatin dual fluorescent and biotin labeling and cell sorting

1-5 x 10^6^ HeLa or HCT116 cells were resuspended in PBS and crosslinked in 1 ml of PBS with 1% formaldehyde. Reaction was stopped using 125 mM glycine. Cells were then permeabilized using 200 μl of 1 x NEBuffer 2 with 1% NP40 for 30 min on ice. 800 ul of 2.5 U of Nt.CviPII, 50 U of DNA Polymerase I and 30 μM of dNTP mix were added to the cells for 2 h at 37°C with constant rotation. dNTP mix contained all four dNTPs plus 6 μM of Fluorescein-12-dATP and 6 μM of biotinylated-dCTP for fluorescent labeling subsequent DNA isolation and sequencing analysis. 50 mM of EDTA were used to block the labeling reaction for 30 min at 37°C. Subsequently, cells were incubated for 15 min at 55°C, spun at 500 g for 5 min at 4°C and resuspended in 1 ml of PBS. 500 μl of PI (Propidium Iodide) /RNase solution was added and incubated with the cells for 20 min at room temperature (Cell Signaling Technology, 4087). The cell suspension was then applied onto FACS tubes with cell strainer cap (Corning, 352235), loaded onto FACS tubes and sorted for G1, S or G2/M phase based on propidium iodide profile using a FACS (Sony, SH800S Cell Sorter). 200,000 of cells sorted in 400 ul of PBS were treated with 40 μl of proteinase K (NEB, P8107S) and 40 μl of 20% SDS and incubated at 65°C for 16 h in order to reverse the DNA crosslinking. Biotin labeled genomic DNA was extracted using the phenol chloroform method.

### NicE-seq library construction

The isolated Biotin-labeled genomic DNA from fixed reactions were sonicated into 150 bp fragments (Covaris) and 200 ng of DNA was mixed with 50 μl of Streptavidin magnetic beads (Invitrogen, 65001), previously blocked using 0.1% cold fish gelatin in 1 × PBS overnight at 4°C in 1 mL of B&W buffer (10 mM Tris-HCl pH 8.0, 1 mM EDTA, 2 M NaCl). Biotin-labeled open chromatin DNA was captured by streptavidin at 4°C for 2 hours with end-over-end rotation. The beads were washed four times with B&W buffer plus 0.05% of Triton X-100 followed by one wash with TE. The beads were resuspended in 50 μl of TE. The DNA was end-repaired, washed twice with B&W buffer plus 0.05% of Triton X-100, dA-tailed and washed with B&W buffer plus 0.05% of Triton X-100. And finally, NEB Illumina adaptor (NEB, E7370S) was ligated and washed twice with B&W buffer plus 0.05% of Triton X-100. A final wash of the bead bound DNA was performed with TE and the bound DNA was resuspended with 20 μls of TE. 10 μl of bound DNA was used for library amplification using PCR (NEB, E7370S). Routinely 8-10 PCR cycles were used to generate enough amount of library DNA for sequencing. The library was examined and quantitated with high-sensitive DNA chip (Agilent, 5067-4627). 4nM was loaded on NextSeq 500 System (Illumina).

### Cell culture, treatment and Western blot

HCT116, HeLa, HeLa-S3, HUT 78, K562, HT1080 and GM12878 cells were grown according to ATCC’s recommendations. HUT 78 cells were treated up to 6 hours with 1 μM of romidepsin (Sigma-Aldrich, SML1175), DMSO being used as a control. Western blot analysis was performed using 5 μg of total cell extract. Antibodies against histone H3 and acetyl-H3 (Lys27) were purchased from Cell Signaling Technology (9715, 8173 respectively). Densitometry analysis were performed using ImageJ. For cell synchronization, HCT116 cells were treated with 2 mM thymidine for 24h, released for 3h followed by addition of 100 ng/mL of nocodazole to the media for 12 h.

### RNA extraction and sequencing

RNA was extracted and purified from HUT 78 cells using Quick-RNA™ Miniprep Kit (Zymo Research, R1054). 1 μg of RNA was used to make libraries for RNA sequencing. Poly(A) mRNA was then isolated using NEBNext^®^ Poly(A) mRNA Magnetic Isolation Module (New England Biolabs, E7490S) and RNA sequencing was performed using NEBNext^®^ Ultra™ II Directional RNA Library Prep Kit for Illumina^®^ (New England Biolabs, E7760S) according to manufacturer’s recommendations. The cDNA libraries were examined and quantitated with high-sensitive DNA chip (Agilent, 5067-4627). 4nM were loaded on NextSeq 500 System (Illumina).

### Bioinformatics analysis

#### Data processing and peak calling

Adaptor and low-quality sequences were trimmed from paired-end sequencing reads using Trim Galore (http://www.bioinformatics.babraham.ac.uk/projects/trim_galore/) with the following setting: --clip_R1 4 --clip_R2 4 --three_prime_clip_R1 4 --three_prime_clip_R2 4. Trimmed read pairs were mapped to the reference genome (human: hg38) using Bowtie2 (37) with the following arguments: --dovetail --no-unal --no-mixed --no-discordant --very-sensitive -I 0 -X 1000. Further, PCR duplicates and mitochondrial reads were removed and only properly aligned read pairs were used for peak calling with MACS2 (38) using ‘macs2 callpeak -f BAMPE -m 4 100 --bdg -SPMR’ options.

#### Cell cycle data analysis

After aligning the raw reads to the reference genome, the technical replicates of G1, S and G2M stages of cell cycle were merged together using samtools merge command (39). Then we downsized the reads from three different stages to the same number of mapped fragments (after excluding PCR duplicates and mitochondrial reads) through random sampling. Then the peaks were called using the same parameter with MACS2 as mentioned above. The Fraction of reads in peaks (FRiP) score was calculated using the deepTools plotEnrichment function (40). Peaks called from G1, S and G2M were compared using the Bedtools (41) and mergepeaks.pl command of Homer (42). First peaks from all the samples are concatenated. Peaks that have at least one base pair overlapping are considered associated and are merged to form a union peak set. Then peaks of individual samples were compared to the union set and were marked as either “unique” or “common”. Last the numbers of “unique” and “common” peaks were summarized from all the samples and were used to make Venn Diagrams in R. Correlation analysis of NicE-seq openchromatin signals were performed with the deeptools plotCorrelation function. Here, affinity-based correlation analysis was performed. The affinity-based method first determines the number of normalized reads that overlap with a set of all the merged peaks from individual samples and then calculates Pearson correlation based on the normalized read count matrix. NicE-Seq peaks were annotated using HOMER annotatePeaks.pl (42). HOMER annotates peaks as promoter (i.e., within 2 kb of known TSS), intergenic, intronic, exon, CpG islands, repetitive elements and other positional categories. After associating peaks with nearby genes and assigning peaks to different genomic features, we also conducted Gene Ontology enrichment analysis using HOMER. Transcription factor binding motifs enrichment near called openchromatin peaks was searched using the HOMER tool findMotifsGenome.pl (42) with default parameters. The enrichment scores (−10log(P-value) of individual TF binding motifs were calculated for G1, S and G2M were summarized into a data matrix in R and a heatmap was then created in to represent stage-specific enrichment of TF binding motifs near the open-chromatin peaks.

TSS of human (hg38) genome were extracted from the NCBI RefGene gene table downloaded from the UCSC Table Browser. HCT116 specific enhancers were downloaded from Enhancer Atlas 2.0 database (43). WGBS data of HCT116 was downloaded from NCBI Gene Expression Omnibus (GEO). ChIP-seq datasets of cell-specific CTCF binding and histone marks (H3K27ac, H3K27me3, H3K4me3) were downloaded human Encyclopedia of DNA Elements (ENCODE) projects and GEO. The TSS, enhancer, WGBS, H3K27ac, H3K27me3, H3K4me3 and CTCF profiles in NicE-vieweq G1, S and G2M was computed with the deeptools computeMatrix and plotheatmap and plotProfile functions (30). ATAC-see data for GM12878 and HT1080 were downloaded from GEO (Supp Table 10). Data processing for HT1080 and GM12878 ATACsee data and HT1080 NicE-viewSeq data was done as described above.

Signal tracks were generated using 100 bp bins using deeptools bamCoverage with the following parameters: --of bigwig --normalizeUsing RPKM

#### HUT 78 Romidepsin treatment

Read processing, alignment, peak calling, peak overlap analysis, correlation analysis and peak annotation was performed as mentioned above.

#### Finding differentially altered peaks

To evaluate the response of HUT 78 to the HDACi romidepsin, we identified the differentially accessible chromatin regions during the drug treatment. For this we created a master peak file by merging the peaks in both control and treated condition from two biological replicates using bedtools merge function as mentioned above. Then the reads corresponding to each peak was evaluated using the multicov module of bedtools and a count matrix was created. Then these read count matrix was normalized and altered peaks were identified using DESeq2 (44) program. Peaks that were showing logFC≥1 at FDR < 0.05 and logFC≤1 at FDR < 0.05 were considered positively altered peaks and negatively altered peaks respectively. These peaks were annotated, to identify their functional relevance, using HOMER as mentioned above. Principal component analysis, clustering and MA plot was generated using DESeq2.

#### RNA-seq analysis

FASTQ files were trimmed using Trim Galore and assessed for quality with FASTQC (https://www.bioinformatics.babraham.ac.uk/projects/fastqc/). None of the samples used in this study were dropped after quality control. Trimmed reads were mapped to hg38 using STAR (45). High-quality mapped reads were then used to estimate transcript abundance using htseq count module (46). Then the count matrix was normalized and differentially expressed genes were identified using the DESeq2 (44). Genes that were showing logFC≥1 at FDR < 0.05 and logFC≤1 at FDR < 0.05 were considered up regulated genes and down regulated respectively. The functional annotation and pathway analysis of the differentially expressed genes were performed using DAVID (47). The differentially expressed genes and genes associated with differentially altered peaks were compared to identify the overlap between NicE-seq and RNA-seq. Spearman correlation was performed between RNA-seq and NicE-seq using deeptools. Differentially expressed genes that are in congruence with NicE-seq peaks were checked for its openness and expression in the gene body and a heatmap was generated. Volcano plot, venn diagram and heatmaps were generated using R.

All datasets used here including their accession numbers are listed in Supp. Table 10.

## Data availability

Nice-viewSeq and RNA seq data performed in this study are available in NCBI Gene Expression Omnibus (GEO) under the accession GSE139253.

## Acknowledgments

We thank W. Jack, C. Carlow and Mihika Pradhan for critical reading of the manuscript, D. Comb, Sir R.J. Roberts and J. Ellard for encouragement. Basic research support for P.O.E., U.S.V., H.G.C., and S.P. was provided by New England Biolabs, Inc.

## AUTHOR CONTRIBUTIONS

S.P conceived and designed the study. H.G.C. and P.O.E performed experiments. U.S.V performed all bioinformatics data analysis. S.P wrote the manuscript with input from all authors. S.P. supervised all aspects of this work.

