## Supplementary figure legends for "Visualization and sequencing of accessible chromatin reveals cell cycle and post romidepsin treatment dynamics"

*Equal contribution

**Supplementary Fig. 1.** 3D NicE-view representation of mitotic cells using LSM880 Airyscan left panel. Right panel represent the profile of mean pixel intensity of DAPI and fluorescein-dATP at an arbitrary diameter of the nuclei.

**Supplementary Fig. 2.** Comparison between accessible chromatin methods with NicE-viewSeq. (A) Pearson correlation of accessible chromatin peak read densities between NicE-viewSeq, NicE-seq and ATAC-seq of HCT116 cells. (B) Representative IGV screen shot of the normalized read density of the NicE-viewSeq, DNase-seq and ATAC-seq libraries of HeLa cells. (C) Pearson correlation of accessible chromatin peak read densities between NicE-viewSeq and ATAC-seq. (D) Genome-wide metagene plot of TSS with ± 2Kb of flanking region of NicE-viewSeq and ATAC-seq analysis.

**Supplementary Fig. 3.** Comparison between NicE-viewSeq and ATAC-see of HT1080 cells. (A) NicE-viewSeq labeling in fixed HT1080 cells using dNTPs supplemented with Texas Red–dATP. Left panel (Texas-Red), middle panel (DAPI) and right panel (merged, DAPI/TexasRed). (B) IGV genomic tracks comparison of accessible chromatin using ATAC-see and NicE-viewSeq. Unfixed indicates native cells, fixed indicates formaldehyde fixed cells. Gene names shown at the bottom. (C) Genome-wide comparison of accessible chromatin between ATAC-see unfixed, ATAC-see fixed and NicE-viewSeq using Pearson correlation. Left panel indicates peak density and right panel read density comparison.

**Supplementary Fig. 4.** Pearson correlation between accessible chromatin region in G1, S and G2M phases using scatter plot display. (A) Accessible chromatin region of HCT116 cell NicE-viewSeq of different cell cycle stages using read counts. (B) Accessible chromatin region of HCT116 cells treated with nocodazole vs G1 and S phase using read counts. (C) Accessible chromatin region of GM12878 cells derived by ATAC-see technology and their correlation analysis in G1, S and G2M phase. For the G1 sequence read density both G1 low and G1 high reads were pulled together. (D) Accessible promoter regions of different cell cycle stages of HCT116 cell using NicE-viewSeq sequence reads.

**Supplementary Fig. 5.** Effect of romidepsin on cells. (A) Quantitative measurement of histone H3K27 acetylation in cells. (B) Hierarchical clustering of two biological replicates of control and romidepsin treated cells. Each set has two technical duplicates as shown. (C) PCA plot showing the variance between the control and romidepsin treated cells. (D) Density plot based on the peak length of open chromatin peaks in control and romidepsin treated cells. (E) Heatmap and signal intensity profile plot of TSS (that includes ± 2Kb of flanking region) in control and after romidepsin treatment. Graphical presentation is shown at the top.

**Supplementary Fig. 6.** RNA transcription post romidepsin treatment and accessible chromatin (A) Hierarchical clustering of three technical replicates of control and romidepsin treated cells for RNA expression analysis. (B) Changes in accessible chromatin regions of repetitive DNA elements. (C) Volcano plot showing the differentially expressed repeat elements post romidepsin treatment (logFC≥1 and FDR<0.05).

**Supplementary Fig. 7.** Gene expression changes post romidepsin treatment. Heatmap of apoptotic related genes showing upregulation post romidepsin treatment.
