## Supplementary figures and images for "Visualization and sequencing of accessible chromatin reveals cell cycle and post romidepsin treatment dynamics"

### Supplementary figures

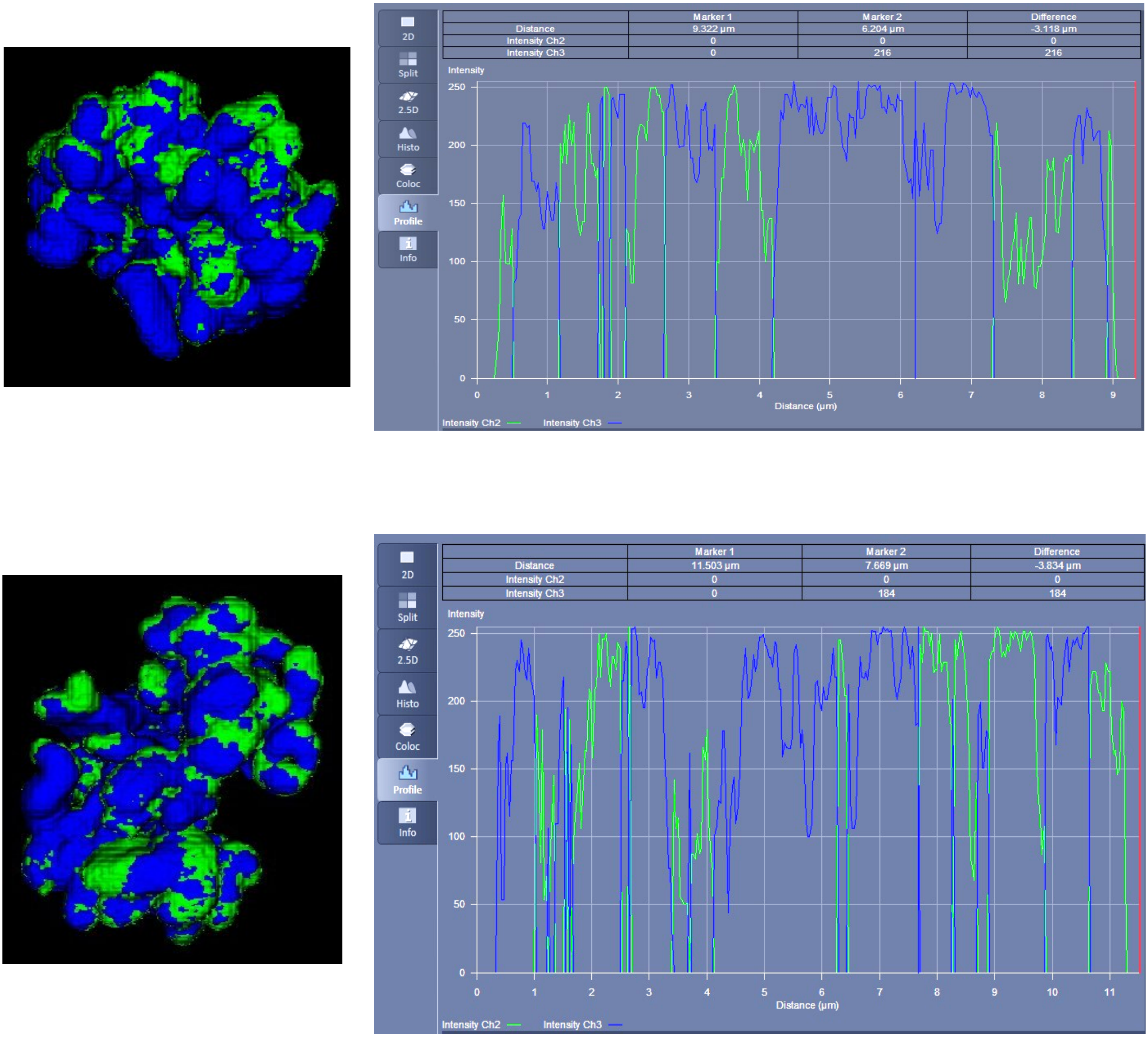

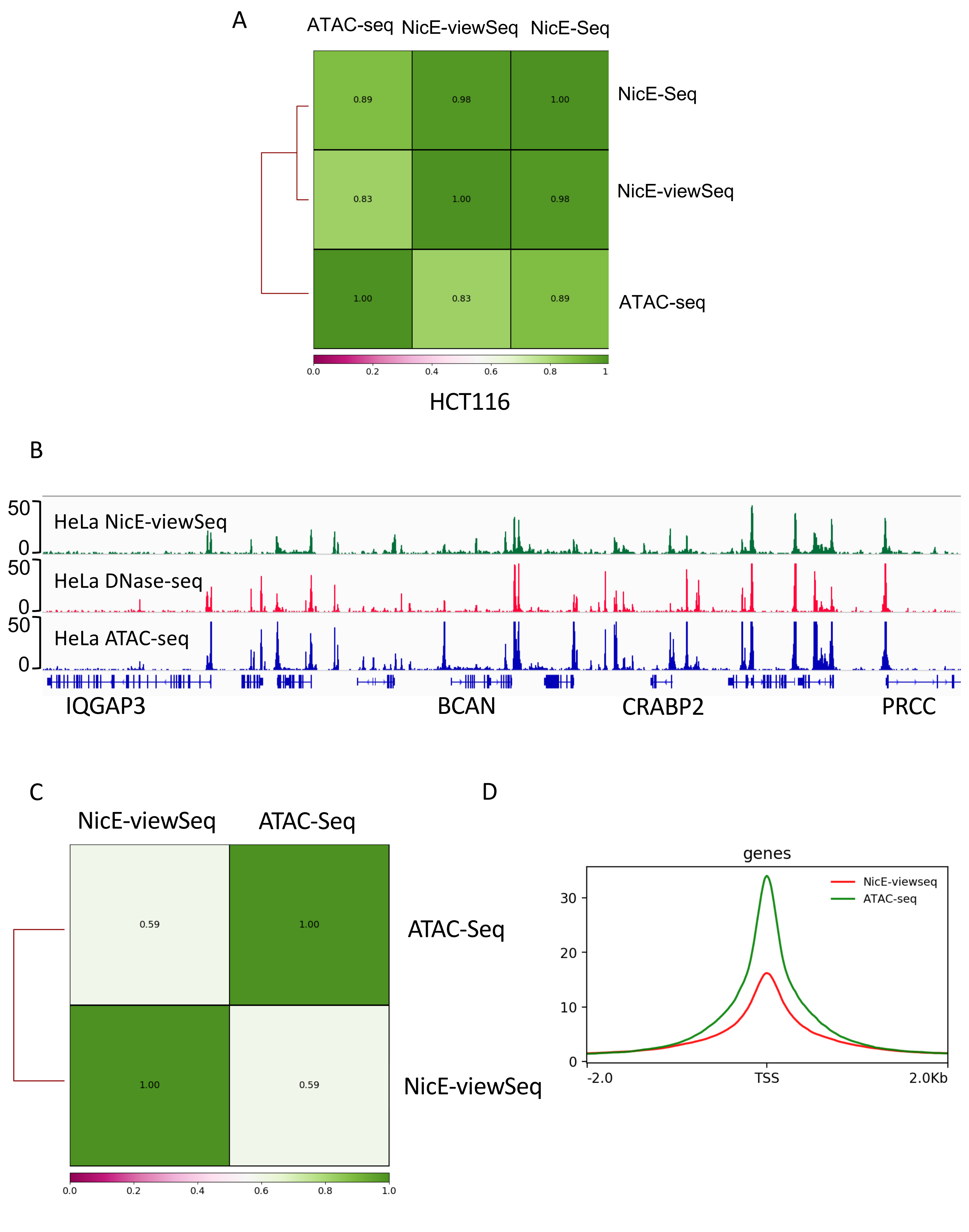

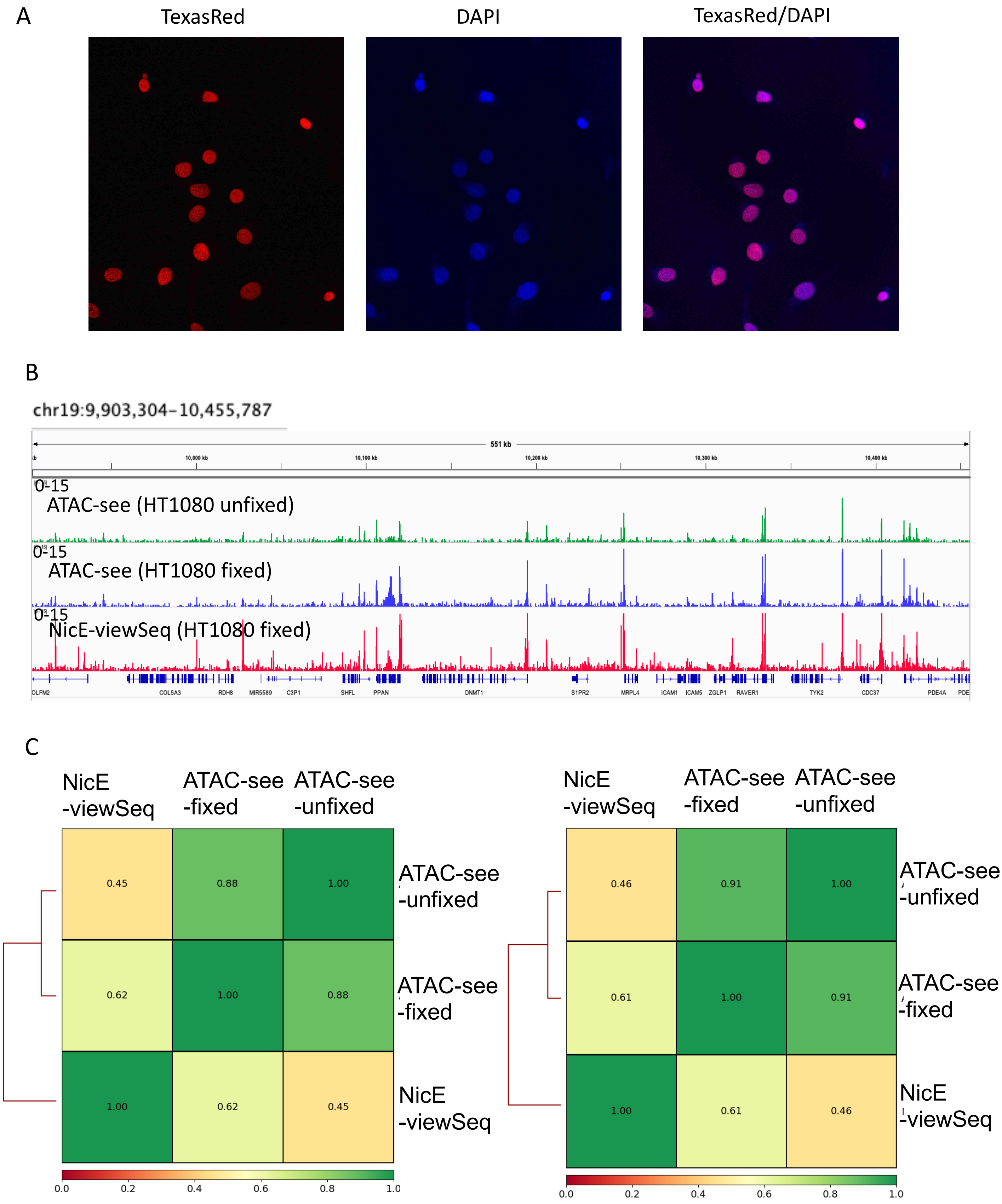

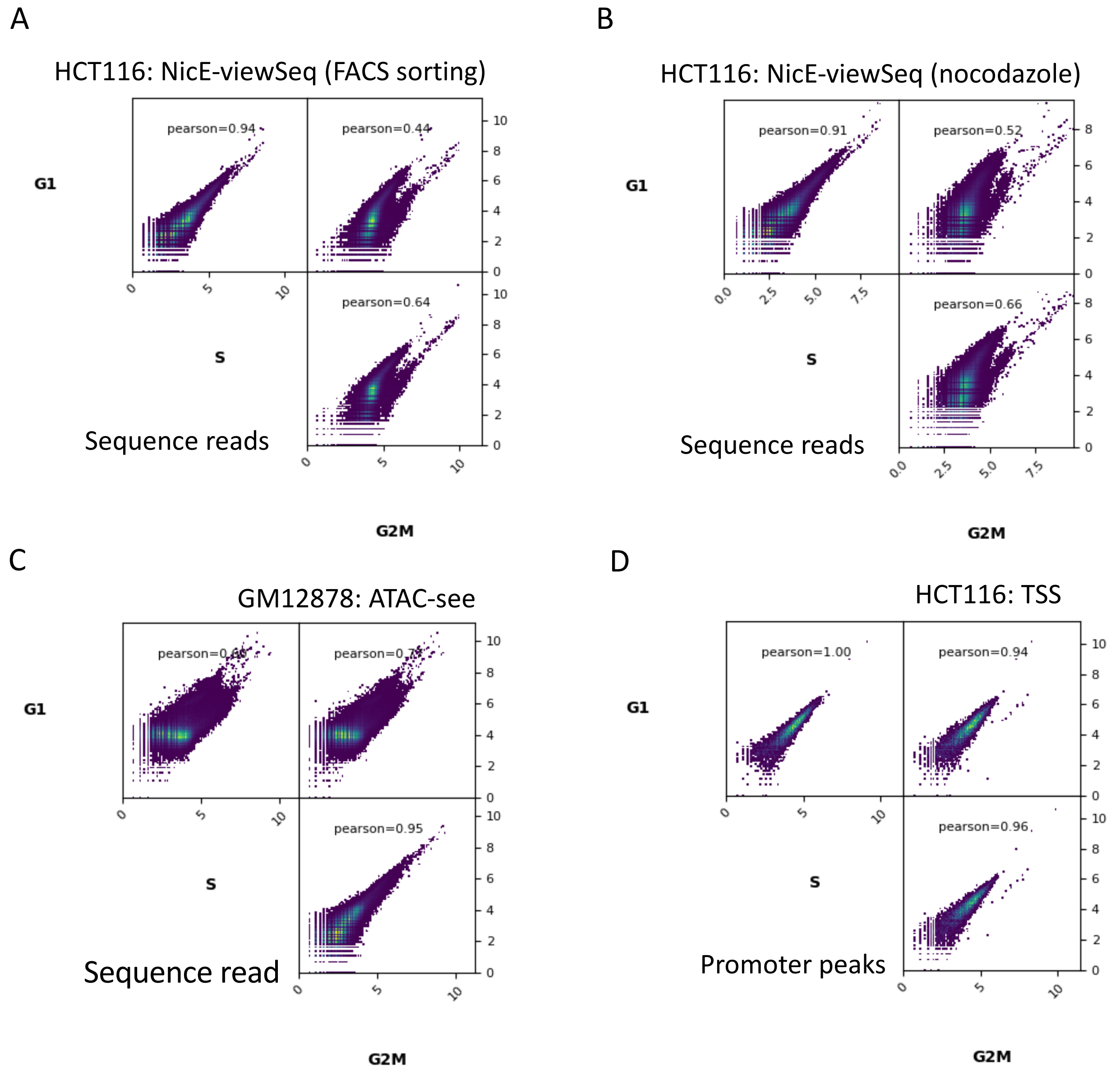

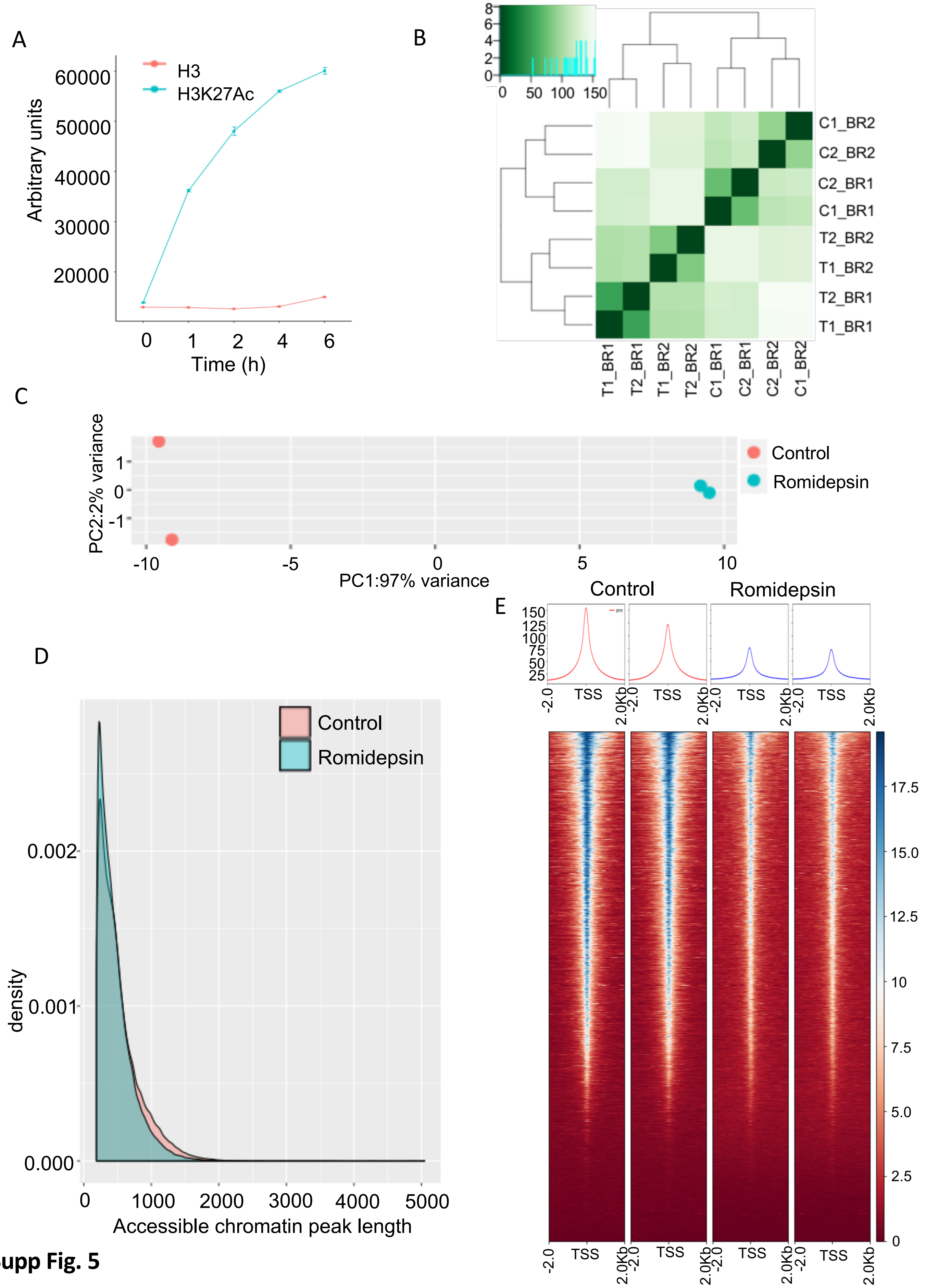

A

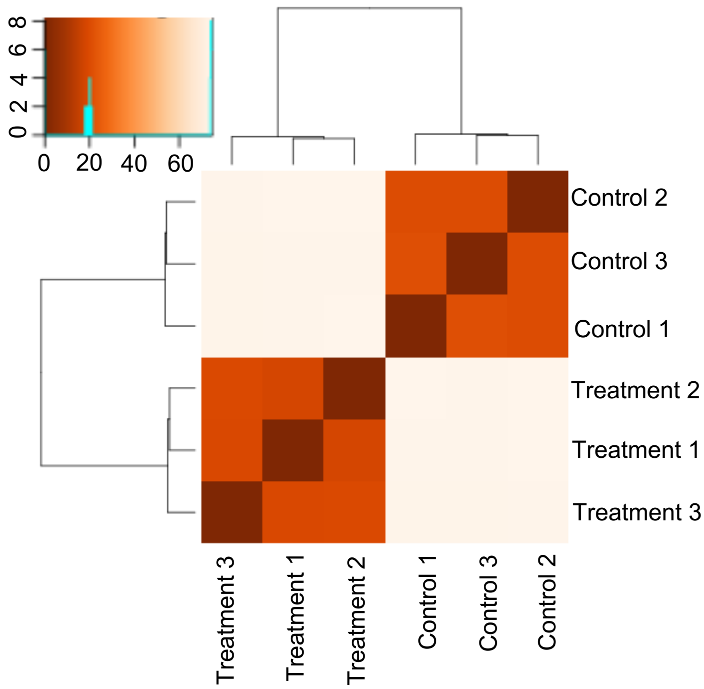


B

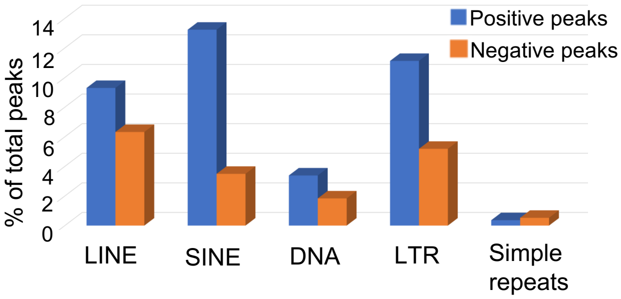


C

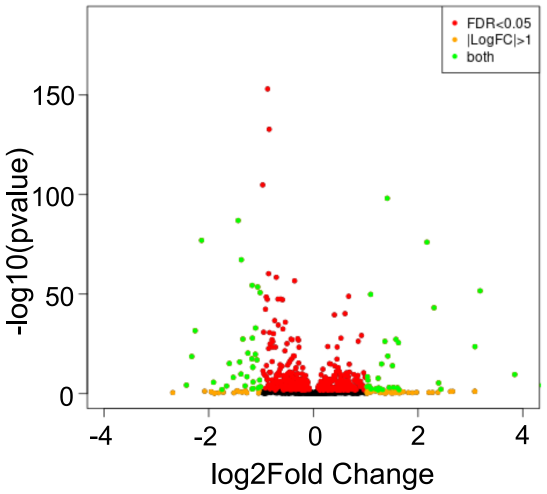


**Supp Fig. 6**

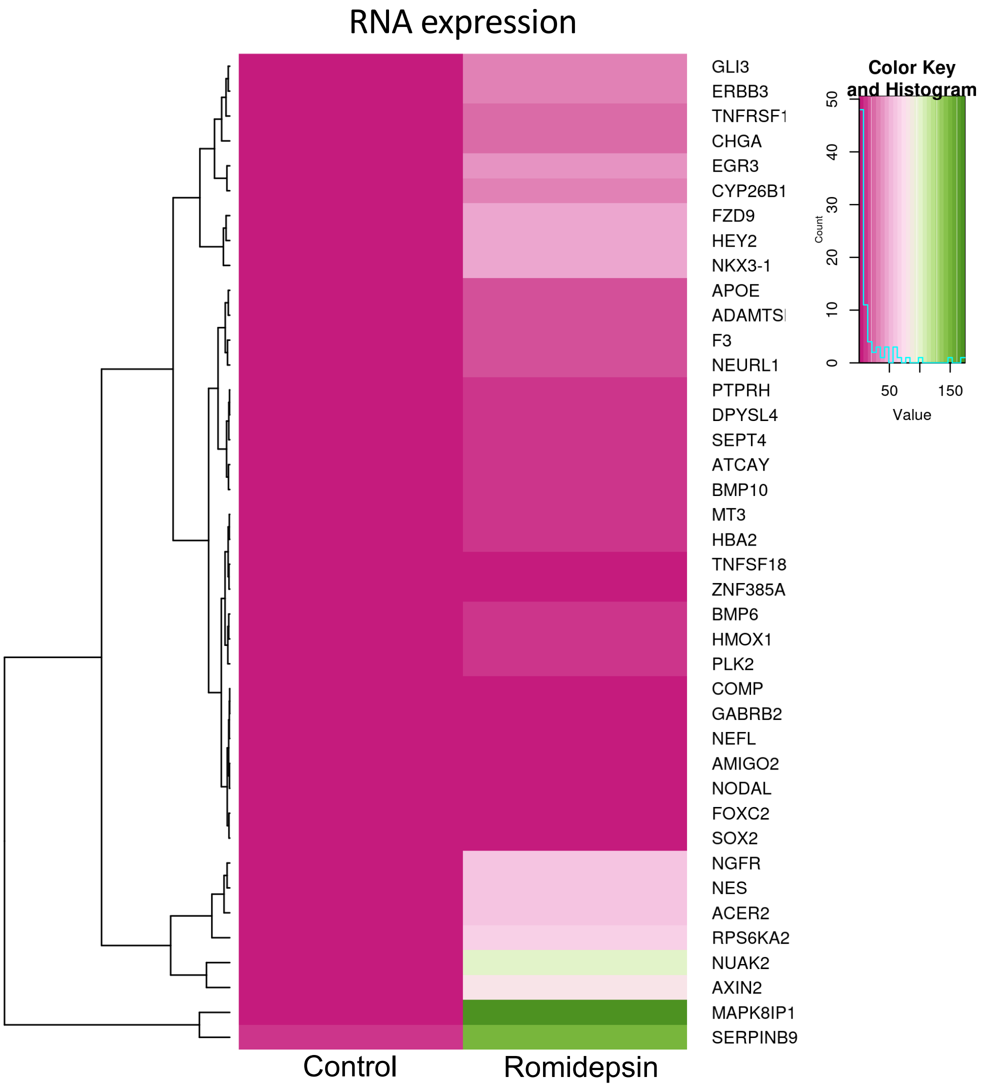
