## Supplementary tables for "Visualization and sequencing of accessible chromatin reveals cell cycle and post romidepsin treatment dynamics"

**Supp Table 2A: HT1080 NicE-viewSeq data statistics**

| Sample Name | Genome | Total reads | Mapped | % mapping rate | Mapped Mt reads | % Mito reads | Reads after mt and dup removal | % duplication | Peaks | Reads in peaks | FRiP | Merged and downsized | Peaks | FRiP |
|---|---|---|---|---|---|---|---|---|---|---|---|---|---|---|
| HT1080_rep1 | hg38 | 94255200 | 91306438 | 96.87 | 13005452 | 14.24 | 74752562 | 0.13 | 155818 | 17691834 | 23.6 | 35000000 | 108187 | 19.4 |
| HT1080_rep2 | hg38 | 134631740 | 130139328 | 96.66 | 19645400 | 15.09 | 104978296 | 0.15 | 191064 | 25499173 | 24.2 | | | |

**Supp Table 2B: HT1080 ATACsee data statistics**

| Sample Name | Genome | Total reads | Mapped | % mapping rate | Mapped Mt reads | % Mito reads | Reads after mt and dup removal | % duplication | Peaks | Reads in peaks | FRiP | Merged and downsized | Peaks | FRiP |
|---|---|---|---|---|---|---|---|---|---|---|---|---|---|---|
| HT1080_fixed_rep1 | hg38 | 56084180 | 54720816 | 97.57 | 35644692 | 65.13 | 15748182 | 0.66 | 58184 | 4262023 | 27.06 | 35000000 | 83725 | 32.07 |
| HT1080_fixed_rep2 | hg38 | 71710552 | 69938308 | 97.53 | 45497728 | 65.05 | 20142234 | 0.67 | 62189 | 5974336 | 29.66 | | | |
| HT1080_unfixed_rep1 | hg38 | 119532370 | 117019648 | 97.9 | 80410160 | 68.71 | 25824714 | 0.74 | 81434 | 7959649 | 30.82 | 35000000 | 89871 | 32.26 |
| HT1080_unfixed_rep2 | hg38 | 102782258 | 100599244 | 97.88 | 71256186 | 70.83 | 20579260 | 0.75 | 66960 | 6009011 | 29.2 | | | |

**p Table 3: HCT116_cell cycle data mapping statistics**

| Sample Name | Genome | Total reads | Mapped | % mapping rate | Mapped Mt reads | % Mito reads | Reads after mt and dup removal | % duplication | Peaks | Reads in peaks | FRiP | Merged and downsized | Peaks | FRiP |
|---|---|---|---|---|---|---|---|---|---|---|---|---|---|---|
| HCT116_G1_rep1 | hg38 | 19593956 | 19377586 | 98.9 | 9678984 | 49.94 | 6906378 | 0.47 | 36229 | 1394711 | 20.19 | 13000000 | 52831 | 24.08 |
| HCT116_G1_rep2 | hg38 | 18922582 | 18709642 | 98.87 | 9537228 | 50.97 | 6567872 | 0.48 | 33753 | 1279350 | 19.48 | | | |
| HCT116_G2M_rep1 | hg38 | 86704740 | 85422142 | 98.52 | 29544684 | 34.58 | 36872662 | 0.49 | 59218 | 9411888 | 25.53 | 13000000 | 26217 | 8.52 |
| HCT116_G2M_rep2 | hg38 | 283476158 | 279234802 | 98.5 | 21988878 | 7.87 | 110941626 | 0.58 | 48313 | 8913706 | 8.03 | | | |
| HCT116_S_rep1 | hg38 | 35121264 | 34705468 | 98.82 | 10413946 | 30.00 | 17537234 | 0.39 | 55016 | 3485713 | 19.88 | 13000000 | 43158 | 17.88 |
| HCT116_S_rep2 | hg38 | 33897862 | 33498776 | 98.82 | 10144770 | 30.28 | 16610220 | 0.4 | 52758 | 3239407 | 19.5 | | | |
| HCT116_Nocodazole_rep1 | hg38 | 32312996 | 31100228 | 96.25 | 3539530 | 11.38 | 26204802 | 0.09 | 26932 | 1802563 | 6.88 | 13000000 | 9924 | 4.02 |
| HCT116_Nocodazole_rep2 | hg38 | 35599714 | 34012490 | 95.54 | 3610446 | 10.61 | 28755798 | 0.09 | 24664 | 1651272 | 5.74 | | | |

**Supp Table 4: HUT78-Romidepsin treatment data mapping statistics**

| Sample Name | Genome | Total reads | Mapped | % mapping rate | Mapped Mt reads | % Mito reads | Reads after mt and dup removal | % duplication | Peaks | Reads in peaks | FRiP |
|---|---|---|---|---|---|---|---|---|---|---|---|
| HUT78_Control_BR1_1 | hg38 | 135344144 | 71903178 | 53.13 | 11062108 | 15.38 | 56263152 | 0.16 | 74067 | 12378946 | 22 |
| HUT78_Control_BR1_2 | hg38 | 56810678 | 32744432 | 57.64 | 2975300 | 9.08 | 28315662 | 0.07 | 48890 | 4627448 | 16.34 |
| HUT78_Rom_BR1_1 | hg38 | 185054838 | 122342734 | 66.11 | 10014348 | 8.18 | 105749474 | 0.1 | 73278 | 13316262 | 12.59 |
| HUT78_Rom_BR1_2 | hg38 | 194516942 | 129106084 | 66.37 | 9685346 | 7.501 | 112118526 | 0.1 | 76997 | 14199984 | 12.67 |
| HUT78_Control_BR2_1 | hg38 | 14785156 | 12906100 | 87.29 | 1103034 | 8.54 | 11071956 | 0.07 | 37873 | 2580143 | 23.30 |
| HUT78_Control_BR2_2 | hg38 | 22784420 | 19898092 | 87.33 | 1759620 | 8.84 | 16823490 | 0.09 | 47252 | 4288773 | 25.49 |
| HUT78_Rom_BR2_1 | hg38 | 30214504 | 27704370 | 91.69 | 1818168 | 6.56 | 24167670 | 0.07 | 41266 | 3122970 | 12.92 |
| HUT78_Rom_BR2_2 | hg38 | 28765638 | 26249012 | 91.25 | 1556658 | 5.93 | 23018474 | 0.07 | 39300 | 3168599 | 13.77 |

**Supp Table 5: HUT78-Romidepsin treatment RNA-seq data mapping statistics**

| Sample Name | Genome | Total reads | Mapped | % mapping rate |
|---|---|---|---|---|
| HUT78_0h_rep1 | hg38 | 14179371 | 13968183 | 98.51 |
| HUT78_0h_rep2 | hg38 | 16021093 | 15779215 | 98.49 |
| HUT78_0h_rep3 | hg38 | 13634246 | 13420157 | 98.43 |
| HUT78_6h_rep1 | hg38 | 24582529 | 24242233 | 98.62 |
| HUT78_6h_rep2 | hg38 | 22702253 | 22396558 | 98.66 |
| HUT78_6h_rep3 | hg38 | 16739670 | 16501536 | 98.51 |

**Supp Table 8: ATAC-see data mapping statistics**

| Sample Name | Genome | Total reads | Mapped | % mapping rate | Mapped Mt reads | % Mito reads | Reads after mt and dup removal | % duplication | Peaks | Reads in peaks | FRiP |
|---|---|---|---|---|---|---|---|---|---|---|---|
| GM12878_G1high_1 | hg38 | 230576648 | 220461192 | 95.61 | 120533192 | 54.67320162 | 10889272 | 0.93 | 12491 | 1442049 | 13.24 |
| GM12878_G1high_2 | hg38 | 278216298 | 267075452 | 96 | 146684238 | 54.92239624 | 9490366 | 0.95 | 10994 | 1236792 | 13.03 |
| GM12878_G1low_1 | hg38 | 190666200 | 181754616 | 95.33 | 108185622 | 59.52290202 | 3039844 | 0.97 | 7386 | 478948 | 15.76 |
| GM12878_G1low_2 | hg38 | 158459444 | 150699740 | 95.1 | 89251878 | 59.22497146 | 2525218 | 0.97 | 6017 | 377464 | 14.95 |
| GM12878_G2_1 | hg38 | 181295586 | 175507114 | 96.81 | 138967168 | 79.18036189 | 6014580 | 0.94 | 14294 | 1126342 | 18.73 |
| GM12878_G2_2 | hg38 | 259593644 | 251934252 | 97.05 | 199269782 | 79.09594683 | 5199090 | 0.96 | 14780 | 1003972 | 19.31 |
| GM12878_S_1 | hg38 | 277182402 | 269962680 | 97.4 | 224370904 | 83.11182272 | 8685688 | 0.94 | 20999 | 1885117 | 21.7 |
| GM12878_S_2 | hg38 | 207235950 | 201302290 | 97.14 | 166933260 | 82.92665722 | 5697082 | 0.94 | 14324 | 1552110 | 17.87 |

**Supp Table 9. GO enrichment of differential peaks**

**A) All differential peaks**

| GO trem | Adjusted p-value |
|---|---|
| Immune response-activating cell surface receptor signaling pathway | 1.68E-33 |
| Intrinsic apoptotic signaling pathway | 6.22E-27 |
| Response to endoplasmic reticulum stress | 1.54E-25 |
| Regulation of defense response to virus | 3.86E-19 |
| Fc-gamma receptor signaling pathway | 8.87E-19 |
| Fc-gamma receptor signaling pathway involved in phagocytosis | 1.71E-18 |
| Fc receptor mediated stimulatory signaling pathway | 1.82E-18 |
| T cell receptor signaling pathway | 2.03E-18 |
| Histone H4 acetylation | 3.59E-18 |
| T cell costimulation | 4.65E-17 |

**B) Positively enriched peaks**

| GO trem | Adjusted p-value |
|---|---|
| Apoptotic signaling pathway | 1.39E-30 |
| Immune response-activating cell surface receptor signaling pathway | 1.04E-25 |
| Intrinsic apoptotic signaling pathway | 2.92E-22 |
| Response to endoplasmic reticulum stress | 6.08E-20 |
| Positive regulation of viral process | 7.20E-18 |
| Positive regulation of nuclease activity | 1.58E-16 |
| Positive regulation of multi-organism process | 2.89E-16 |
| Activation of signaling protein activity involved in unfolded protein response | 4.49E-16 |
| Regulation of defense response to virus | 4.82E-16 |
| Antigen processing and presentation of peptide antigen | 6.78E-15 |

**C) Negatively enriched peaks**

| GO trem | Adjusted p-value |
|---|---|
| T cell activation | 6.78E-13 |
| B cell activation | 4.28E-09 |
| Positive regulation of myeloid cell differentiation | 6.19E-08 |
| Regulation of alpha-beta T cell proliferation | 2.47E-07 |
| Lymphocyte activation involved in immune response | 5.80E-07 |
| B cell differentiation | 8.12E-07 |
| Positive regulation of leukocyte differentiation | 7.95E-07 |
| T cell receptor signaling pathway | 3.656E-06 |
| Leukocyte activation involved in immune response | 4.8843E-06 |
| Positive regulation of myeloid leukocyte differentiation | 5.3098E-06 |

**Supp Table 10: Data sets used in this study**

| Cell line | Condition/Antibody and Replicate number | Experiment | Single End (SE) or Paired End (PE) | Genome | Source | Accession |
|---|---|---|---|---|---|---|
| HCT116 | G1-1 | NicE-viewSeq | PE | hg38 | This study | GSE139253 |
| HCT116 | G1-2 | NicE-viewSeq | PE | hg38 | This study | GSE139253 |
| HCT116 | S-1 | NicE-viewSeq | PE | hg38 | This study | GSE139253 |
| HCT116 | S-2 | NicE-viewSeq | PE | hg38 | This study | GSE139253 |
| HCT116 | G2M-1 | NicE-viewSeq | PE | hg38 | This study | GSE139253 |
| HCT116 | G2M-2 | NicE-viewSeq | PE | hg38 | This study | GSE139253 |
| HCT116 | G2M-1_Nocodazole_treated | NicE-viewSeq | PE | hg38 | This study | GSE139253 |
| HCT116 | G2M-2_Nocodazole_treated | NicE-viewSeq | PE | hg38 | This study | GSE139253 |
| HeLa | G1-1 | NicE-viewSeq | PE | hg38 | This study | GSE139253 |
| HeLa | G2M-1 | NicE-viewSeq | PE | hg38 | This study | GSE139253 |
| HUT78 | Control-BR1_1 | NicE-viewSeq | PE | hg38 | This study | GSE139253 |
| HUT78 | Control-BR1_2 | NicE-viewSeq | PE | hg38 | This study | GSE139253 |
| HUT78 | Romidepsin_BR1_1 | NicE-viewSeq | PE | hg38 | This study | GSE139253 |
| HUT78 | Romidepsin_BR1_2 | NicE-viewSeq | PE | hg38 | This study | GSE139253 |
| HUT78 | Control-BR2_1 | NicE-viewSeq | PE | hg38 | This study | GSE139253 |
| HUT78 | Control-BR2_2 | NicE-viewSeq | PE | hg38 | This study | GSE139253 |
| HUT78 | Romidepsin_BR2_1 | NicE-viewSeq | PE | hg38 | This study | GSE139253 |
| HUT78 | Romidepsin_BR2_2 | NicE-viewSeq | PE | hg38 | This study | GSE139253 |
| HUT78 | Control-1 | RNA-seq | PE | hg38 | This study | GSE139253 |
| HUT78 | Control- 2 | RNA-seq | PE | hg38 | This study | GSE139253 |
| HUT78 | Control- 3 | RNA-seq | PE | hg38 | This study | GSE139253 |
| HUT78 | Romidepsin-1 | RNA-seq | PE | hg38 | This study | GSE139253 |
| HUT78 | Romidepsin-2 | RNA-seq | PE | hg38 | This study | GSE139253 |
| HUT78 | Romidepsin-3 | RNA-seq | PE | hg38 | This study | GSE139253 |
| HT1080 | Rep1 | NicE-viewSeq | PE | hg38 | This study | GSE139253 |
| HT1080 | Rep2 | NicE-viewSeq | PE | hg38 | This study | GSE139253 |

| | | | | | | |
|---|---|---|---|---|---|---|
| HCT116 | CTCF | ChIP-seq | SE | hg38 | Gertz et al., 2013 | ENCSR000BSE |
| HCT116 | H3K27Ac | ChIP-seq | SE/PE | hg38 | ENCODE project consortium, 2012 | ENCSR661KMA |
| HCT116 | H3K4me3 | ChIP-seq | SE/PE | hg38 | ENCODE project consortium, 2012 | ENCSR333OPW |
| HCT116 | H3K27me3 | ChIP-seq | SE/PE | hg38 | ENCODE project consortium, 2012 | ENCSR810BDB |
| HCT116 | Control-1 | WGBS | PE | hg38 | Ponnaluri et a.,2017 | GSE97889 |
| HCT116 | Control-2 | WGBS | PE | hg38 | Ponnaluri et a.,2017 | GSE97889 |
| HeLa | Control | ATAC-seq | PE | hg38 | Cho et al.,2018 | GSE106145 |
| HCT116 | Control | ATAC-seq | PE | hg38 | Cho et al.,2018 | GSE97583 |
| HeLa | Control | DNase-seq | PE | hg38 | ENCODE project consortium, 2012 | ENCSR959ZXU |
| HCT116 | Control | DNase-seq | SE | hg38 | Maurano et al., 2012 | ENCSR000ENM |
| GM12878 | G1_high_1 | ATAC-see | PE | hg38 | Chen et al., 2016 | GSE79921 |
| GM12878 | G1_high_2 | ATAC-see | PE | hg38 | Chen et al., 2016 | GSE79921 |

| | | | | | | |
|---|---|---|---|---|---|---|
| GM12878 | G1_low_1 | ATAC-see | PE | hg38 | Chen et al., 2016 | GSE79921 |
| GM12878 | G1_low_2 | ATAC-see | PE | hg38 | Chen et al., 2016 | GSE79921 |
| GM12878 | S_1 | ATAC-see | PE | hg38 | Chen et al., 2016 | GSE79921 |
| GM12878 | S_2 | ATAC-see | PE | hg38 | Chen et al., 2016 | GSE79921 |
| GM12878 | G2_1 | ATAC-see | PE | hg38 | Chen et al., 2016 | GSE79921 |
| GM12878 | G2_2 | ATAC-see | PE | hg38 | Chen et al., 2016 | GSE79921 |
| HT1080 | Unfixed_rep1 | ATAC-see | PE | hg38 | Chen et al., 2016 | GSE76006 |
| HT1080 | Unfixed_rep2 | ATAC-see | PE | hg38 | Chen et al., 2016 | GSE76006 |
| HT1080 | fixed_rep1 | ATAC-see | PE | hg38 | Chen et al., 2016 | GSE76006 |
| HT1080 | fixed_rep2 | ATAC-see | PE | hg38 | Chen et al., 2016 | GSE76006 |
